# Gene circuit performance characterization and resource usage in a cell-free ‘breadboard’

**DOI:** 10.1101/000885

**Authors:** Dan Siegal-Gaskins, Zoltan A. Tuza, Jongmin Kim, Vincent Noireaux, Richard M. Murray

**Affiliations:** Division of Biology and Biological Engineering, California Institute of Technology, Pasadena, CA, USA; Faculty of Information Technology, Pazmany Peter Catholic University, Budapest, Hungary; School of Physics and Astronomy, University of Minnesota, Minneapolis, MN, USA; Department of Control and Dynamical Systems, California Institute of Technology, Pasadena, CA, USA

**Author notes:** These authors contributed equally to this work.

**Keywords:** cell-free systems, biological circuit prototyping, crosstalk, *in vitro* synthetic biology, RNA aptamer

## Abstract

The many successes of synthetic biology have come in a manner largely different from those in other engineering disciplines; in particular, without well-characterized and simplified prototyping environments to play a role analogous to wind-tunnels in aerodynamics and breadboards in electrical engineering. However, as the complexity of synthetic circuits increases, the benefits—in cost savings and design cycle time—of a more traditional engineering approach can be significant. We have recently developed an *in vitro* ‘breadboard’ prototyping platform based on *E. coli* cell extract that allows biocircuits to operate in an environment considerably simpler than but functionally similar to *in vivo*. The simplicity of this system makes it a promising tool for rapid biocircuit design and testing, as well as for probing fundamental aspects of gene circuit operation normally masked by cellular complexity. In this work we characterize the cell-free breadboard using real-time and simultaneous measurements of transcriptional and translational activities of a small set of reporter genes and a transcriptional activation cascade. We determine the effects of promoter strength, gene concentration, and nucleoside triphosphate concentration on biocircuit properties, and we isolate the specific contributions of essential biomolecular resources—core RNA polymerase and ribosomes—to overall performance. Importantly, we show how limits on resources, particularly those involved in translation, are manifested as reduced expression in the presence of orthogonal genes that serve as additional loads on the system.

## ABBREVIATIONS

TX: transcription
TL: translation
MGapt: malachite green RNA aptamer
UTR: untranslated region
RNAP: RNA polymerase
NTP: nucleoside triphosphate
RBS: ribosome binding site

## 1 Introduction

The field of synthetic biology has matured to the point where biological parts are regularly assembled into modestly complex circuits with wide-ranging applications (1). Unfortunately, the development of new biological circuits typically involves long and costly *ad hoc* design cycles characterized by trial-and-error and lacking the prototyping stage essential to other engineering disciplines. More often than not, designed circuits fail to operate as expected. The reason for these failures is in many cases related to *context*: the poorly characterized environment in which the system is operating (2–4). This includes the finite and variable (from cell to cell, condition to condition, and time to time) pools of biomolecular resources such as transcription/translation machinery and nucleoside triphosphates (NTPs), weak control over the component DNA concentrations, unpredicted interactions between components and circuits and their cellular hosts (5, 6), and any number of other system properties with unknown or unknowable effects.

We have recently developed an *in vitro* biomolecular ‘breadboard’ based on *E. coli* cell extract that provides a functional environment similar to *in vivo* but with significantly reduced complexity (7, 8). DNA and mRNA endogenous to the cells is eliminated during extract preparation, so that transcription–translation circuits of interest may be operated in isolation without interference by a cellular host. The cell-free breadboard also allows for a degree of control over reaction conditions and component concentrations that cannot be achieved *in vivo*. As a prototyping platform, the cell-free breadboard provides for a considerable reduction in circuit design cycle time, not only because of its relative simplicity when compared with *in vivo*, but also because it eliminates much of the lengthy cloning and cell transformation steps typically required in biocircuit development (9, 10). Indeed, cell-free applications for synthetic biology are quickly expanding (11, 12). But beyond its potential as an improved circuit development platform, our cell-free breadboard has another significant advantage: its simplicity reveals important details of biocircuit operation normally masked by cellular complexity.

In this work we show a detailed and quantitative characterization of the cell-free breadboard—an essential precursor to any biocircuit development and testing application—and explore a number of fundamental aspects of biocircuit operation not easily studied *in vivo*. Central to our work is the use of a novel reporter that combines the malachite green RNA aptamer and a fluorescent protein for a real-time and simultaneous read-out of the system’s transcription and translation activity. We establish the functional implications of intrinsic biocircuit properties such as component concentration and promoter strength, as well as those of the extrinsic biomolecular resource pool that includes nucleoside triphosphates (NTPs) and transcription/translation machinery. Finally, through a systematic characterization of the effect of loads on transcriptional and translational performance, we show how limits on essential resources, particularly those involved in translation, manifest themselves in the form of reduced expression and ‘crosstalk’ between orthogonal genes. Implications for biocircuit prototyping are discussed.

## 2 Results and discussion

### 2.1 **A combined transcription–translation reporter**

To best characterize transcription– and translation–level performance in the cell-free breadboard platform, we use a reporter construct encoding a green or cyan fluorescent protein (deGFP/deCFP) along with the malachite green RNA aptamer (MGapt) in the 3′ untranslated region (UTR) (Fig. 1A). These fluorescent proteins have been previously designed for maximal expression in the cell-free system when transcribed from *E. coli* promoters (7). The 35-base MGapt sequence contains a binding pocket for the malachite green dye (13) and allows mRNA dynamics to be easily monitored in cell extract with high temporal resolution—as compared to, e.g., radio-labeling and gel analysis (14)—and over a wide dynamic range; concentrations of purified deGFP-MGapt transcript as low as 37.5 nM and as high as 3 *µ*M were effectively detected using the MGapt fluorescence signal (Fig. S1). (In principle, the upper limit for RNA detection is determined by the amount of malachite green dye.)

**Figure 1:**
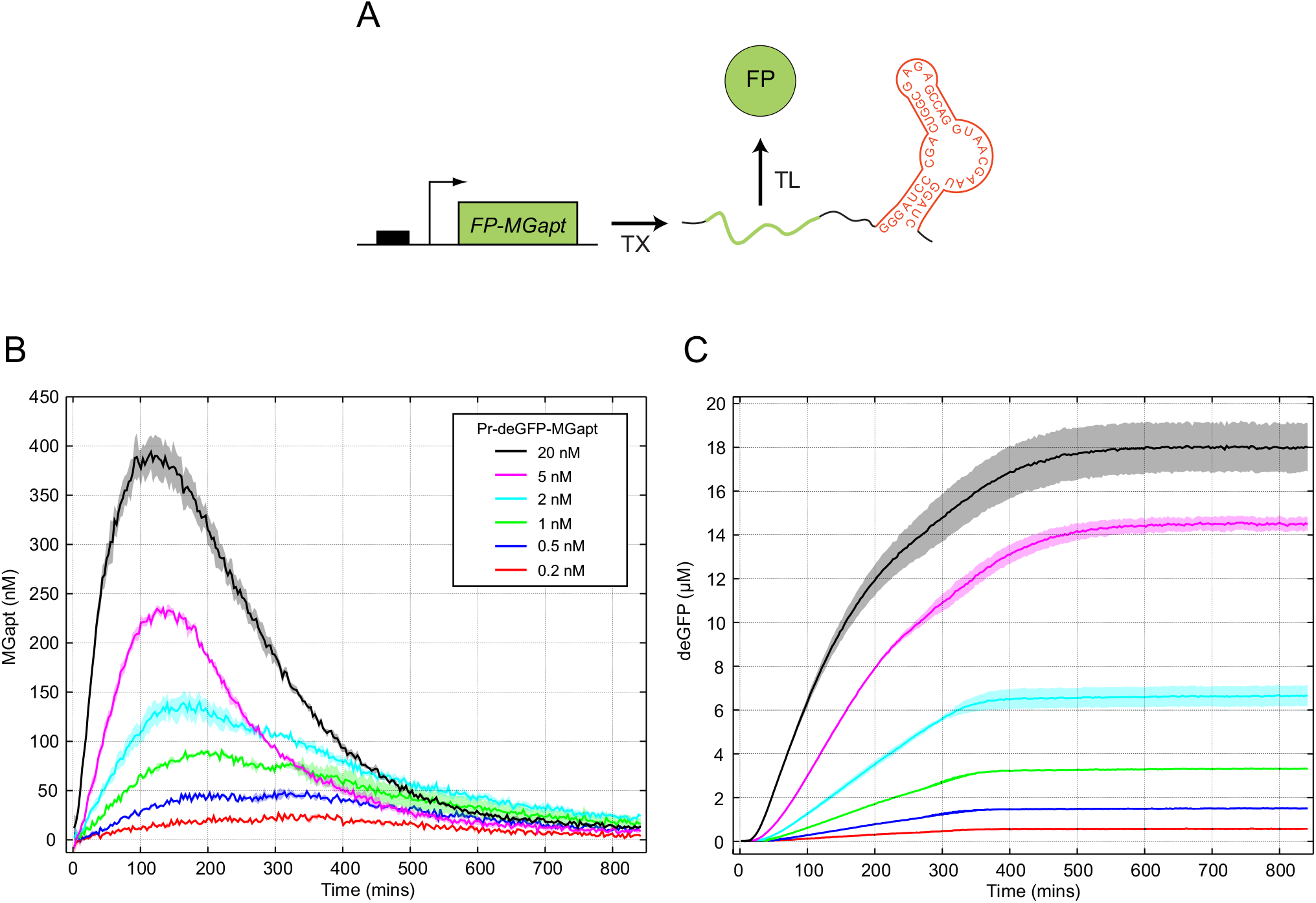
A combined transcription–translation reporter reveals basic features of our cell-free breadboard system. (A) The reporter construct encodes an optimized fluorescent protein (FP) along with the malachite green RNA aptamer (MGapt) in the 3′ UTR. (B) Transcription kinetics reported by MGapt for six different template concentrations. Shaded regions indicate standard error over three replicates. (C) Translation kinetics reported by deGFP for six different template concentrations. Endpoint values represent the total amount of active fluorescent protein produced.

Measurements of the decay of deGFP-MGapt transcripts in the cell-free breadboard show that degradation is well-described by single exponential decay with 16–18 min half-lives for a wide range of concentrations (Fig. S2A). The decay curves for high concentrations of deGFP-MGapt (2 and 3 *µ*M) follow those of the lower concentrations only to 15 min, after which the half-lives increase dramatically (Fig. S2B). This result is consistent with multimer formation occuring in the highly concentrated purified RNA stock (12 *µ*M); the nucleic acid folding program NUPACK (15) predicts 20% dimer formation for 4 *µ*M deGFP-MGapt mRNA and 60% dimer formation for 12 *µ*M deGFP-MGapt mRNA at 37 ^◦^C.

The MGapt fluorescence signal is well-separated from those produced by commonly-used green, yellow, and cyan fluorescent proteins. Furthermore, the inclusion of MGapt in the 3′ UTR of deGFP has little effect on final deGFP levels (Fig. S3), and expression kinetics reported with MGapt are consistent with real-time PCR measurements (Fig. S4). Recent studies also have indicated that MGapt is compatible with *in vivo* characterization (16). Taken together these results suggest that deGFP-MGapt is a reliable tool for concurrent measurement of transcription and translation activity and a strong addition to the set of similar tools used for real-time monitoring of biocircuit performance (cf., (17, 18)).

### 2.2 **Constitutive gene expression under standard conditions**

In the ‘ideal’ system with unlimited resources and conditions unchanging with time, the dynamics of MGapt and deGFP may be described by a set of ordinary differential equations:

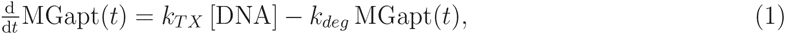

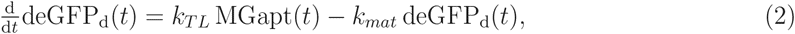

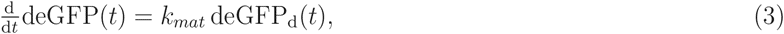

with constants *k_T X_*, *k_deg_*, *k_TL_* and *k_mat_* and template concentration [DNA], and with deGFP_d_ representing the ‘dark’ (immature) deGFP. (There is no protein degradation term in the model as the necessary degradation machinery is absent from the standard cell-free breadboard.) Models of this type are common and have the advantage that they may be solved exactly; for example, this simple model predicts a steady-state MGapt concentration (= *k_T X_*[DNA]*/k_deg_*) and a rate of deGFP production that approaches a constant (= [DNA]*k_T X_k_T L_/k_deg_*) when *k_mat_t*, *k_deg_t* ≫ 1. However, when the operational demands of a biocircuit outstrip the available resources, or those resources exhibit a time-dependent activity, simple models such as this provide little guidance. Precisely how resource limits manifest themselves is the focus of the remainder of this work, with the ideal model used as a point of comparison.

With deGFP-MGapt under the control of a strong constitutive promoter, Pr, and 5′ UTR containing a strong ribosome binding site (RBS), the MGapt expression profiles show clear signs of transcriptional resource limits (Fig. 1B). Saturation of the transcription machinery occurs as template DNA concentration increases; for sufficiently early times, the rates of MGapt production are well described by:

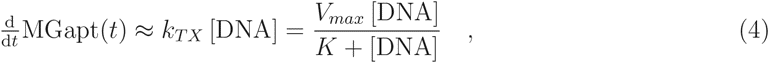

with *K* ∼ 15 nM and *V_max_* ∼ 90 nM min^-1^ (Fig. S5). RNA production does not continue indefinitely, as evidenced by the absence of a steady-state level of MGapt and, at late times, kinetics that follow a pure exponential decay (Fig. S6A). We find that the MGapt degradation rate constant is not the same for all template concentrations, but rather increases with increasing DNA concentrations (Fig. S6B). Between 2 and 5 nM DNA template, there is a distinct qualitative change in the MGapt profiles, from curves characterized by relatively broad peaks and slow decays to ones that are more sharply-peaked.

The deGFP expression profiles are shown in Fig. 1C. As with the MGapt curves, the time during which protein is produced is limited; however, the precise time *t_end,TL_* past which no additional protein is produced is different above and below the 2–5 nM DNA threshold. For all concentrations below 2–5 nM, *t_end,TL_* appears fixed at ≈340 minutes. At high concentrations, *t_end,TL_* is substantially later (at ≈500 minutes). These results suggest that the cessation of protein production is not likely to be purely due to consumption of resources by the transcription–translation machinery. Similar results have been noted previously (19) with the suggestion that a number of other processes, including NTP hydrolysis and enzyme denaturation, may lead to early termination of protein synthesis reactions (20, 21); however, it is worth noting that in our cell-free breadboard the protein production time is considerably longer than is typical for other batch-mode cell-free reactions (19, 22).

The deGFP production rates (while *t < t_end,TL_*) also show differences between the high and low concentration regimes. The rates could be expected to correlate with the level of MGapt since, in the ideal system described by Eqs. (1)–(3), we have that

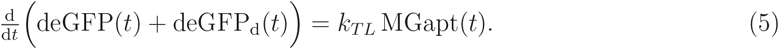

Indeed, in the first hour of expression, the time derivative of the (spline fit) deGFP concentration is proportional to the MGapt concentration (Fig. S7A). At later times, the behavior of the system is no longer ideal and 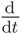deMGapt(*t*) and MGapt(*t*) diverge, although there remain significant qualitative similarities between them (Fig. S7B). One difference between deGFP production rates and MGapt levels is apparent at low DNA concentrations, where the rate of deGFP production is approximately constant for times ∼90 mins *< t <* ∼300 mins despite the fact that MGapt is not. There is no similar period with a constant deGFP production rate above 5 nM DNA.

We use two simple metrics to assess system performance under different conditions: the complete time integral of MGapt concentration (*∫*MGapt) and the endpoint concentration of deGFP ([deGFP]*_end_*). These particular metrics were chosen because they reflect the total transcription– and translation–level capacity of the system and are easily comparable between experimental runs. For the ideal system we have that:

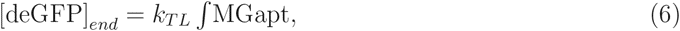

as may be seen by integrating Eq. (5) over the full length of the experimental run. (Here we have used the fact that the endpoint concentration of immature deGFP is relatively small for late times, i.e., [deGFP]_d_*_,end_* ≪ [deGFP]_*end*_.) In the cell-free breadboard, a proportionality relation exists only for the first hour of expression below the 2–5 nM DNA threshold, i.e., deGFP(*t\**) ∝ 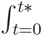 MGapt(*t*)d*t* for *t*\* ≤ 60 mins and [DNA] ≤ 2 nM (Fig. 2A). At higher template concentrations and later times, performance deviates from ideality as the chemical make-up of the system is substantially changed and the effects of resource limits are more pronounced (Fig. S8). However, it is at this point when *∫*MGapt and [deGFP]*_end_* are no longer simply related by Eq. (6) that they may separately provide insights into how specific resource limitations are manifested.

**Figure 2:**
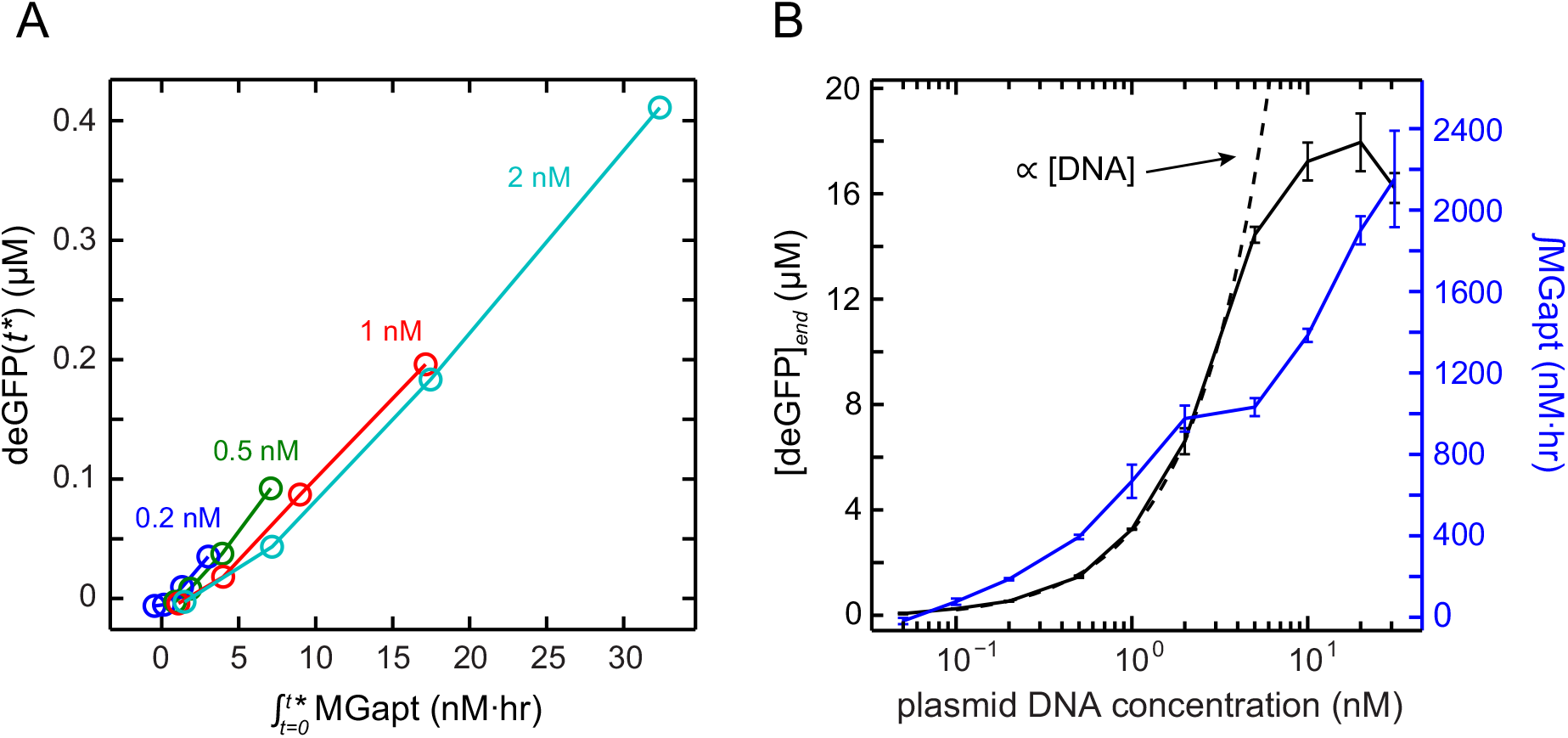
(A) deGFP at various times *t*\* versus MGapt concentration integrated from time *t* = 0 to *t* = *t*\* for *t*\* = 15, 30, 45, and 60 minutes and a range of DNA template concentrations. (B) Endpoint deGFP and integrated MGapt as a function of DNA template concentration. The endpoint deGFP level is proportional to the amount of template up to ∼2–5 nM.

Plotting *∫*MGapt and [deGFP]*_end_* as a function of plasmid concentration (Fig. 2B), we clearly see the ‘linear’ regime in which [deGFP]*_end_* is proportional to DNA concentration and a ‘saturation’ regime in which [deGFP]*_end_* versus DNA concentration is sublinear. The existence of a linear regime may be easily predicted from the ideal model Eq. (6):

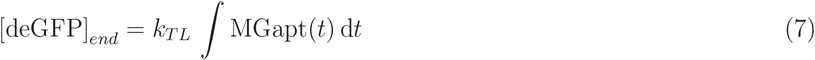

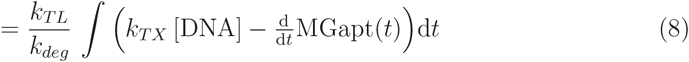

which, in the limit that *k_T__X_* [DNA] ≫ 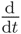MGapt(*t*) reduces to the simple relation

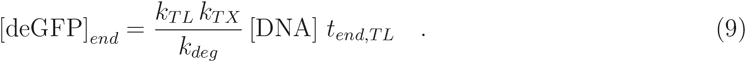

The linear and saturation regimes correspond to the qualitative differences in the MGapt and deGFP expression curves described above. Surprisingly, we see no significant change in *∫*MGapt at the regime transition point.

### 2.3 **Performance as a function of promoter strength and DNA concentration**

To establish how promoter strength affects transcription and translation in the cell-free breadboard, we tested the reporter construct under the control of two additional constitutive promoters Pr1 and Pr2 made weaker than Pr by single base mutations in the -35 and -10 region, respectively (see Materials and Methods). The concentration at which the system transitions from the linear regime to the saturation regime is increased for these weaker promoters, up to ∼10 nM for Pr1 and ∼20 nM for Pr2 (Fig. S9), consistent with other recent work suggesting a maximum capacity between 16 and 32 nM DNA (9). Thus, we see a performance trade-off between DNA concentration and promoter strength: a weaker promoter allows for linear regime performance with higher template concentrations.

The relationship between DNA template concentration, promoter strength, integrated RNA, and final protein concentration is not a simple one (Fig. 3A). For example, with 2 nM DNA, *∫*MGapt produced using Pr1 and Pr2 is 40% and 15% of the Pr value, respectively, and [deGFP]*_end_* is 11% and 0.4% of that produced by Pr. The percentages are different with 20 nM DNA: *∫*MGapt produced using Pr1 and Pr2 is 70% and 32% of Pr, respectively, and [deGFP]*_end_* is 30% and 2% of Pr. (Separate comparisons of *∫*MGapt and [deGFP]*_end_* for the three promoters can be found in Fig. S10.)

**Figure 3:**
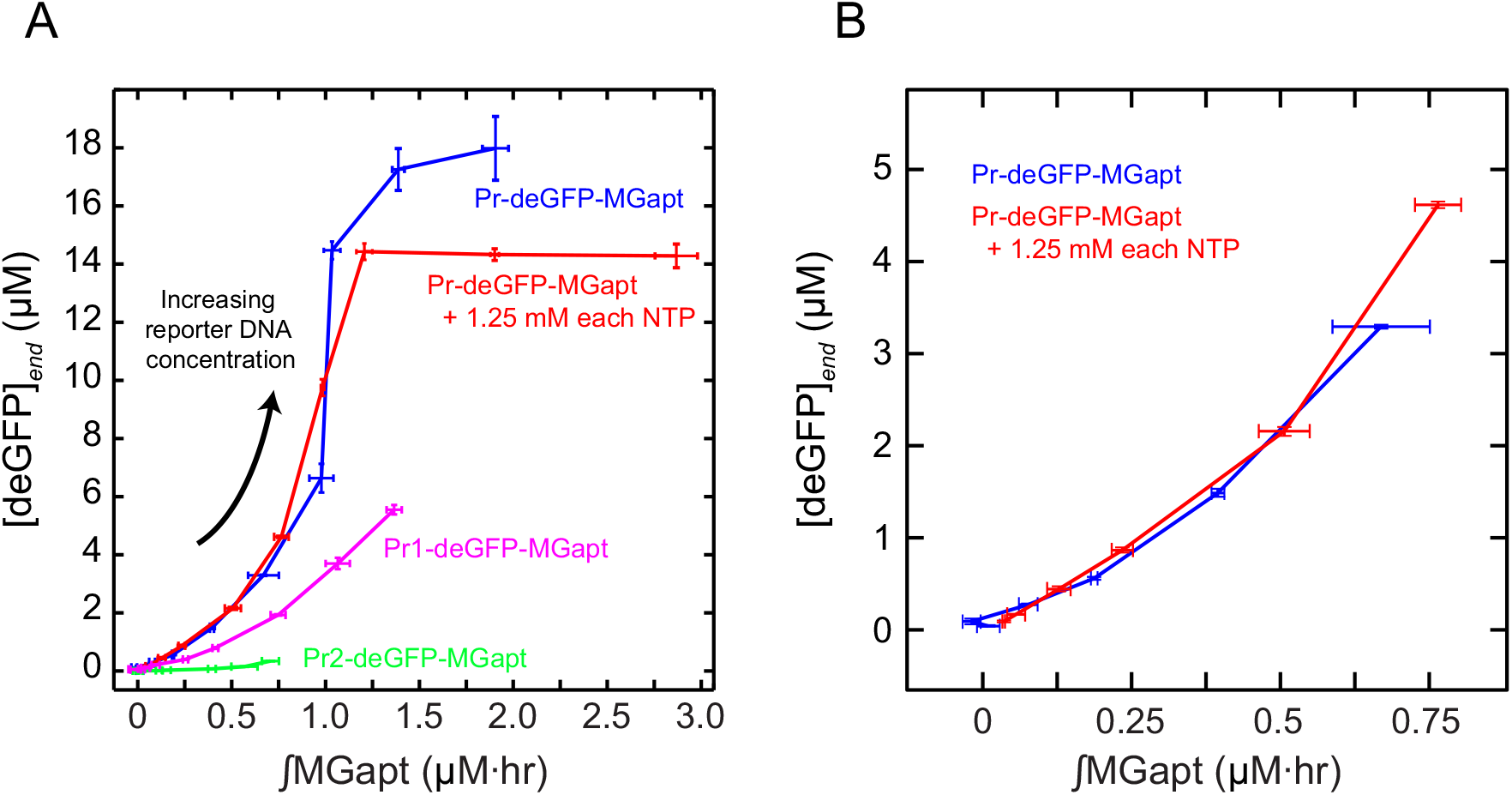
(A) Endpoint deGFP versus integrated MGapt for three different promoters (Pr, Pr1, and Pr2) and with the cell-free breadboard supplemented with 1.25 mM of each of the four NTPs. DNA template concentrations are 0.02–20 nM. (B) Comparison of linear regime performance, with and without additional NTPs.

The dramatic increase in the Pr promoter [deGFP]*_end_*–*∫*MGapt curve found at the regime transition point may be explained by the differential transcriptional dynamics in the linear and saturation regimes—distributed versus peaked—coupled with decreasing activity of the translation machinery. A consequence of a time-dependent reduction in translational efficiency, as reported for other cell-free systems (19), is that although transcription may take place late in the experiment, the resulting mRNAs are less translatable.

### 2.4 **The role of NTPs**

The standard platform contains the natural NTPs essential for biocircuit operation, in concentrations of 1.5 mM ATP and GTP and 0.9 mM CTP and UTP (23). Among their many cellular functions, ATP and GTP play a crucial role in translation, and all four NTPs are used in transcription as substrates in the synthesis of RNA. NTPs thus serve to couple a biocircuit’s transcription and translation layers together, with an impact that is not intuitively obvious but that can be significant (24). As a result, understanding precisely how changes in NTP concentration affect performance is of paramount importance.

We supplemented the system with an additional 1.25 mM of each NTP, an increase of ∼83% ATP/GTP and ∼138% CTP/UTP. In the high DNA concentration (saturation) regime, the additional NTPs support a considerable increase in *∫*MGapt but lead to a reduction in [deGFP]*_end_* of up to 20% (Fig. 3A), a result of a dramatic broadening of the MGapt expression curves and compression of deGFP curves at late times (Fig. S11). This suggests that while NTPs may help at the transcription level, the excess transcripts are not translatable, and that perhaps the resources used to produce those transcripts may have been taken at the expense of reporter protein production.

In the low DNA concentration (linear) regime, additional NTPs lead to an ∼40–50% increase in [deGFP]*_end_* (Fig. 3B). This may be primarily attributed to an increase in the value of *t_end,TL_* (to ∼450–500 minutes; see Fig. S11). That is, the rate of production is relatively fixed but the productive period is extended. *∫*MGapt also increases at low DNA concentrations. Save for the increase in *t_end,TL_*, the extra NTPs have little effect on the shapes of the MGapt and deGFP profiles (Fig. S11).

### 2.5 **Performance of a simple transcription–translation cascade**

We also investigated the effect of adding an intermediate layer of transcription and translation on our reporters. The two-stage ‘cascade’ circuit consists of constitutively-expressed T7 RNAP under the control of Pr, Pr1, or Pr2 and the deGFP-MGapt construct downstream of a T7-specific promoter (Fig. 4). (T7 RNAP is convenient to use for this purpose since, unlike the native *E. coli* RNAP, it is a single-subunit RNAP that is easy to incorporate onto a single plasmid and it does not compete with the core RNAP for sigma factors.) If the consumption of NTPs by transcription/translation is in fact performance-limiting, we would expect the output of the cascade to be reduced relative to constitutive expression since more NTPs are used in its operation. Alternatively, if expression is limited (at least in part) by a reduction in the activity of the native RNAP, the introduction of T7 RNAP may extend the lifetime of the system.

**Figure 4:**
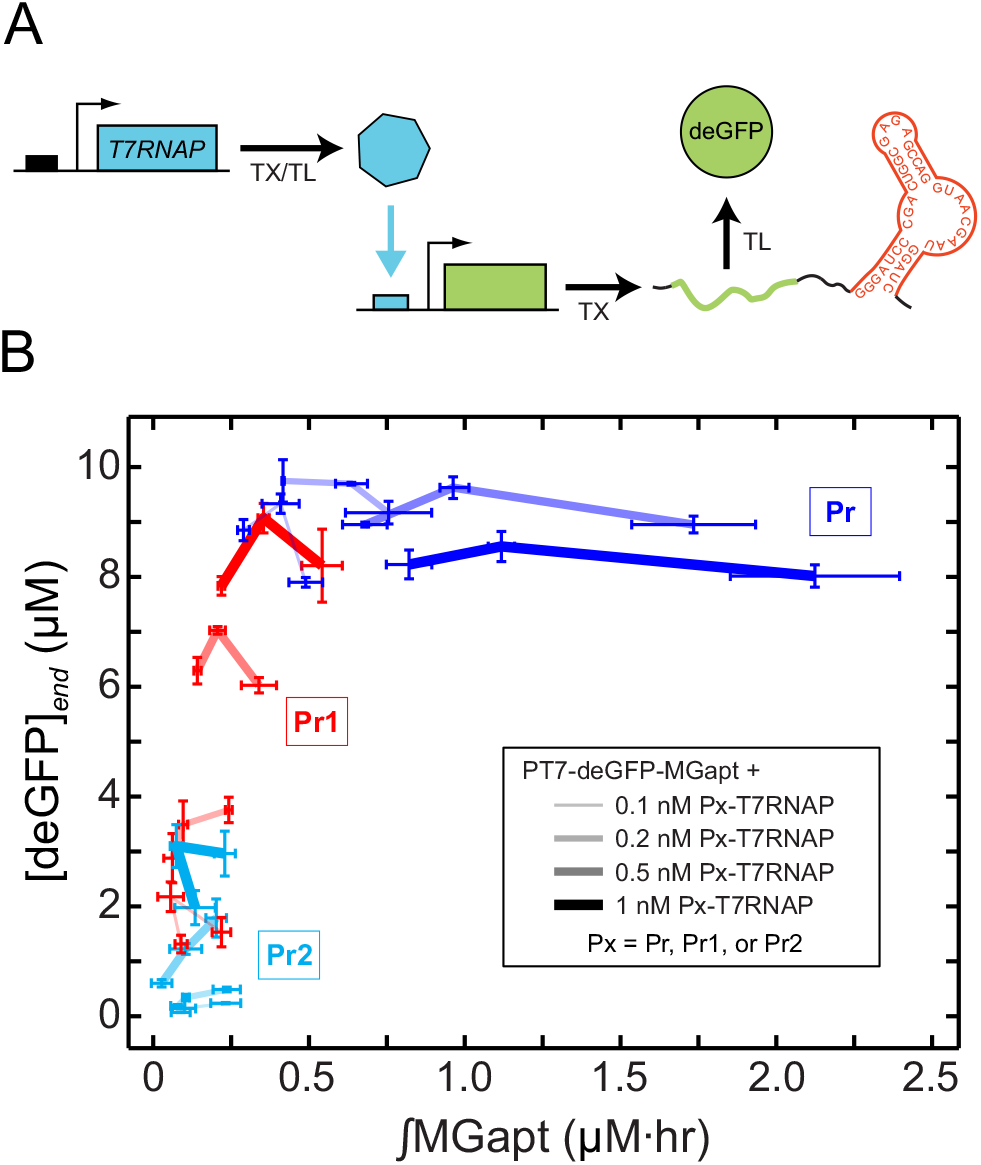
(A) The T7 cascade consists of constitutively-expressed T7 RNAP under the control of Pr, Pr1, or Pr2 and the deGFP-MGapt construct downstream of a T7-specific promoter. (B) Endpoint deGFP versus integrated MGapt for the T7 cascade with Pr-, Pr1-, or Pr2-T7 RNAP upstream of the PT7-deGFP-MGapt reporter (1, 2, and 10 nM). There exist regions of overlap where high concentrations of weaker first-stage promoters perform similarly to low concentrations of stronger first-stage promoters.

There are substantial qualitative differences in T7 cascade expression as compared with a single-stage constitutive promoter. RNA increases rapidly and exhibits a long, slow decay (Fig. S12), and there is a ∼30-60 minute delay in protein expression (Fig. S13). A higher reporter DNA concentration leads to a shorter delay and faster rise in expression, but the final deGFP concentration is often below the level achieved with a lower reporter concentration. This suggests a trade-off in cascade-driven protein production that may be the result of fuel consumption: if deGFP is produced more quickly, then the production appears to arrest sooner.

The cascade protein output is largely determined by the strength of the promoter that drives T7 RNAP production (Fig. 4). Weaker promoters (Pr1 and Pr2) lead to a wide range of deGFP levels with only small variations in the Pr1- and Pr2-T7 RNAP plasmid concentrations, and for any fixed concentration of the T7 RNAP plasmid, changes in reporter concentration (over an order of magnitude) do not affect deGFP output appreciably. When the strong Pr promoter is used, deGFP levels saturate at a level independent of the Pr-T7 RNAP concentration while MGapt levels vary substantially. The T7 cascade thus provides for independent tuning of RNA and protein outputs. There exist regions of overlap where cascades with high concentrations of weaker first-stage promoters behave identically to low concentrations of stronger first-stage promoters; for example, 1 nM Pr1-T7 RNAP produces an output ([deGFP]*_end_* and *∫*MGapt) similar to that produced by 0.1 nM Pr-T7 RNAP, and 1 nM Pr2-T7 RNAP lies between 0.1 and 0.2 nM Pr1-T7 RNAP. This equivalence was not present with the one-stage simple expression and may be due to the fact that in all versions of the cascade the promoters driving deGFP are identical.

In the simple expression case we found that adding NTPs to the system led to a considerable increase in transcription in the saturation regime, but that the excess transcripts were not translated. We set out to see how the same addition of NTPs affects output of the T7 cascade with a strong first-stage promoter. As before, we supplemented the system with an additional 1.25 mM of each NTP. The resulting kinetics can be seen in Figs. S14 and S15, and compared with Figs. S12 and S13. Again we see a significant increase in the transcriptional activity; peaks are taller and broadened and the differences between different PT7-deGFP-MGapt concentrations are more pronounced. And while we do not see a decrease in deGFP as we did with the Pr-deGFP-MGapt construct, there is little to be gained at the translational level by supplying excess NTPs.

### 2.6 **Resource utilization and crosstalk between orthogonal genes**

Recent studies have revealed that unintended and/or indirect coupling between genes, or ‘crosstalk’, can arise as a side-effect of resource sharing (25–31). Resources include those which are consumed during circuit operation (e.g., NTPs) as well as those like the transcription/translation machinery and other enzymes that are not consumed but are limited and shared across different parts of the circuit. To clearly distinguish between crosstalk at the transcription stage (that may arise due to competition for RNAP or housekeeping sigma factor) and at the translation stage (arising from, e.g., a limited ribosome pool), we used a deCFP-MGapt reporter construct to assay system performance in the presence of two different ‘loads’: (1) the ‘native’ deGFP, containing the same UTR with strong RBS used throughout this work, and (2) deGFP with the RBS deleted from the UTR (ΔRBS-deGFP). (We use the term ‘load’ to refer to components that consume some of the fixed transcriptional and translational capacity of the system, in analogy with electric loads that consume electric power.) Reporter and loads are all placed under the control of the same Pr promoter. The loads are orthogonal to the reporter in the sense that they have no direct regulatory interaction with it, e.g., as activating or repressing transcription factors. With this particular setup, crosstalk at the transcription and translation levels appears as changes in MGapt and deCFP fluorescence signals, respectively. The use of the Pr-ΔRBS-deGFP construct guarantees that any crosstalk is strictly transcriptional, as no RBS is present to sequester ribosomes away from the production of deCFP. In principle, purely translational loads such as high concentrations of purified mRNA could be used; however, purified deGFP-MGapt transcripts at high concentration exhibit nonideal behavior (as noted earlier; cf. Fig. S2), and the effect of this nonideality on reporter and load output is unknown.

We find that an increase in loading generally leads to a decrease in reporter expression, although the effect on transcription is different from that on translation (Fig. 5). For example, a similar ∼250 nM·hr variation in *∫*MGapt is observed for all reporter concentrations as loading increases. At the translation level, the effect is strongly concentration-dependent: the influence of the load on [deCFP]*_end_* is small at 1 nM reporter DNA but significant at 10 nM reporter. Thus, as the amount of reporter DNA increases, the demand for resources required for production of deCFP increases as well and the translational crosstalk becomes much more pronounced. This result underscores the fact that there is a maximum translation capacity limiting the total amount of protein that can be produced; as [deGFP]*_end_* goes up, [deCFP]*_end_* necessarily goes down. The top right corners of the plots in Fig. 5 represent a performance regime that appears to be inaccessible.

**Figure 5:**
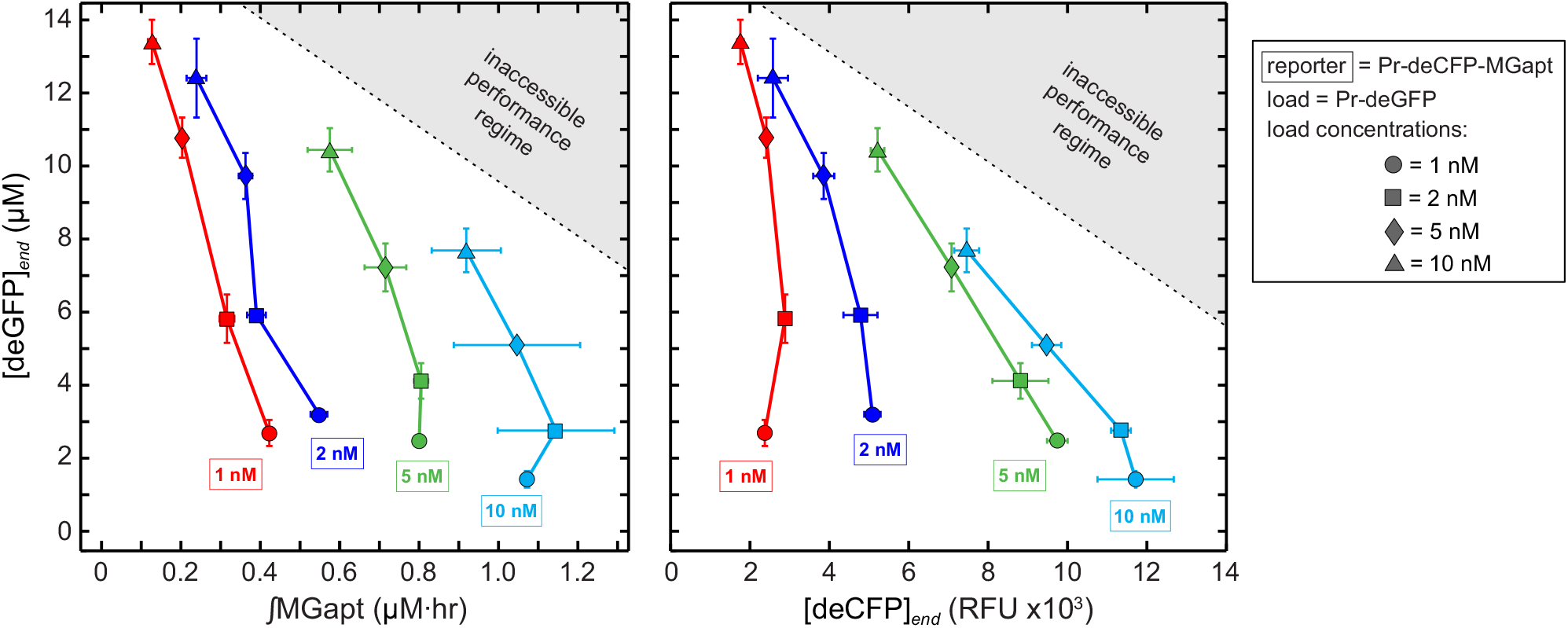
Effect of expression of load Pr-deGFP on concurrent expression of reporter Pr-deCFP-MGapt. Loading is quantified as endpoint deGFP concentration and plotted against integrated MGapt (left) and endpoint deCFP (right). Variation in *∫*MGapt with increasing load is similar for all reporter concentrations, whereas at the translation level the crosstalk effect is highly dependent on load and reporter concentrations. There is a maximum translation capacity to the system that limits the total amount of protein that can be produced, as indicated by the inaccessible performance regime.

The loading effects seen in Fig. 5 suggest that translation resources may be more limiting to system performance. To confirm this result, we compare the *∫*MGapt–[deCFP]*_end_* relationships for Pr-deGFP and Pr-ΔRBS-deGFP (Fig. 6). When the RBS is present, we see that in general an increase in load leads to a decrease in *∫*MGapt and [deCFP]*_end_* (Fig. 6, left). When the RBS is absent (Fig. 6, right), for the most part the load has no effect on performance, except for at high concentrations of both load and reporter, at which point we note a decrease in [deCFP]*_end_* and increase in *∫*MGapt.

**Figure 6:**
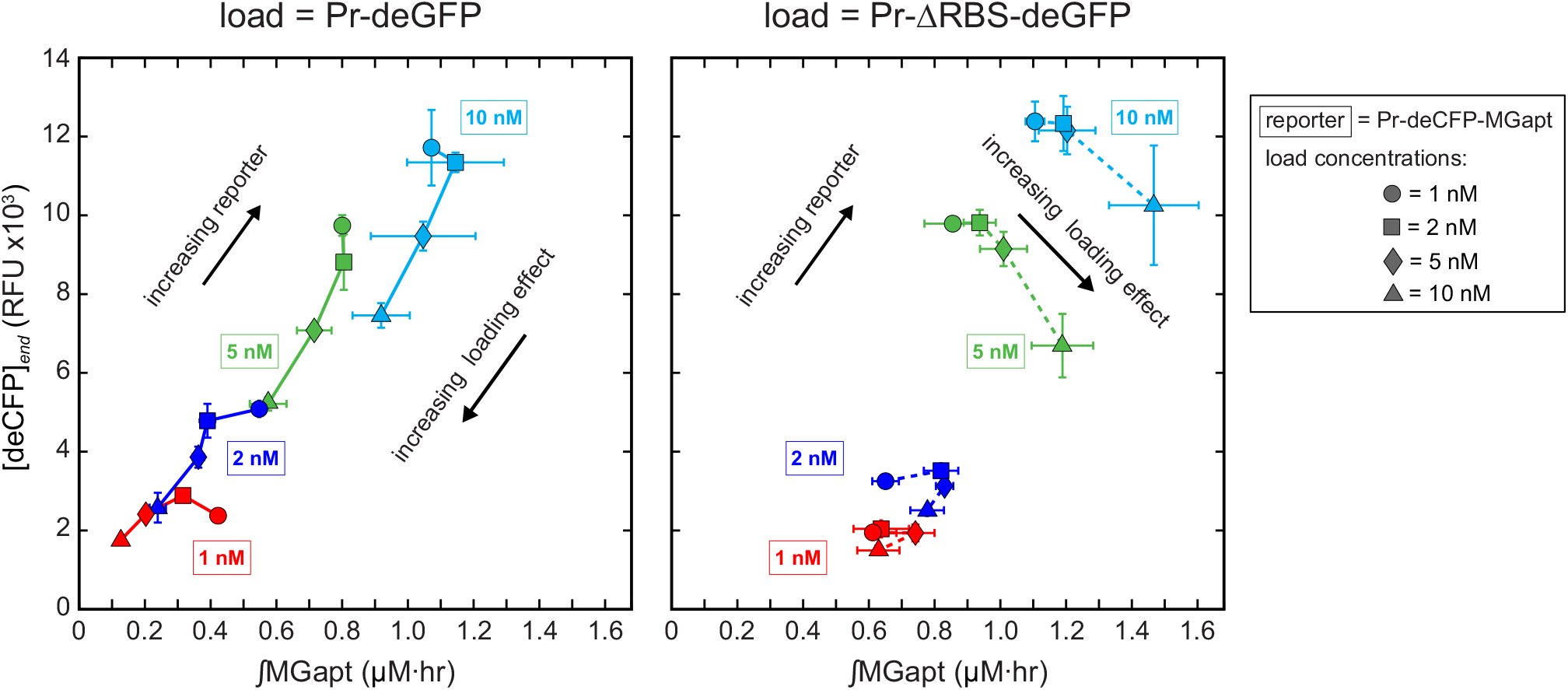
Effect of expression of loads Pr-deGFP (left) and Pr-ΔRBS-deGFP (right) on concurrent expression of reporter Pr-deCFP-MGapt. Increasing Pr-deGFP tends to decrease both *∫*MGapt and [deCFP]*_end_*. Without the RBS, only when load and reporter concentrations are high do we see a crosstalk effect: a decrease in [deCFP]*_end_* and increase in *∫*MGapt.

### 2.7 Relevance for *in vivo*

While it can be expected that specific circuit behaviors will manifest themselves to different degrees in cell-free environments versus *in vivo*, nevertheless there is significant potential for cell-free work to contribute to understanding how circuits function in living systems. One example may be found in our ‘resource competition’ assays, through which we were able to quickly and clearly observe the translation machinery serving as a significant limiting resource. It has been suggested—by recent theoretical work (29) as well as by several other studies on ribosome utilization (30, 32–34)—that similar ribosome loading effects exist *in vivo*, despite the fact that live cells are able to produce additional translation machinery.

Other consequences of a limited ribosome pool found in the cell-free breadboard may also be found *in vivo*. In particular, it is known that translating ribosomes protect their template mRNAs from the action of endonucleases (35) and that ribosome spacing is a determinant of degradation rate (36). Thus, if demands on a system are such that the available ribosome pool is insufficient to densely cover the number of transcripts, increased endonucleolytic activity would lead to an increase in the RNA degradation rate constant. Consistent with this hypothesis we note an increase in the degradation rate constant when the system transitions from the linear to the saturation regime (Fig. S6).

Ribosomes are not, of course, the only molecular resource that may be found in short supply and that can, as a result of their limited number, lead to crosstalk between unrelated circuits. Examples include the *E. coli* ClpXP protein degradation machinery (26) and transcription factors (TFs) with relatively large numbers of targets (37). (Although the latter theoretical study dealt specifically with TF titration, the thermodynamic model may be generalizable to other ‘targeting’ biomolecules.) One particularly interesting class of candidate sources of crosstalk is the RNases. It has been suggested that competition for a relatively small number of RNases by a large number of RNA molecules can introduce unintended correlations (24), and evidence for this may be found in our results: the addition of the untranslated ΔRBS-deGFP load in amounts higher than or comparable to the reporter results in an increase in *∫*MGapt (Fig. 6). In this case, the load presents a large number of new targets for degradation enzymes, drawing them away from the RNA reporter and thus indirectly leading to the increase in *∫*MGapt. We are currently unaware of *in vivo* results that demonstrate crosstalk via RNases; however, given the ribosome loading effects seen in both cell-free and *in vivo* systems, it is an intriguing possibility worthy of exploration, with particular relevance for RNA-based synthetic regulatory circuits (38–40).

The cell-free breadboard may be useful in predicting or confirming a number of other *in vivo* effects arising due to resource limits. For example, in a recent modeling study it was suggested that different combinations of promoter and RBS strengths can result in comparable protein output with different loads on the cellular expression machinery, and that codon usage can introduce a bottleneck that impacts the expression of other genes (41). The degree of precise control that exists in the cell-free breadboard—for example, control over DNA concentration and known induction levels without an intervening cell membrane—makes it an ideal platform for investigating this and other related questions.

### 2.8 **On biocircuit prototyping**

Despite recent developments in standardized part libraries and rapid assembly tools (e.g., (42, 43)), synthetic biology still lacks the accepted prototyping platforms and protocols common to other engineering disciplines. For the purposes of prototyping, one particular advantage of our cell-free breadboard is the rapid testing cycle it permits: save for the initial cloning, transformation, and plasmid preparation, none of the individual assays performed in this work required the many hours of cell treatment typically needed for *in vivo* studies. With plasmids in hand, the time from cell-free experiment setup to first results is a matter of minutes.

Problems associated with limits on the cell-free breadboard system capacity may be mitigated when operating in regimes that yield predictable response; for example, in the low DNA concentration regime where the protein production rate is approximately constant until a well-defined, concentration-indepedent end time, and the final protein concentration is proportional to the amount of template DNA (Fig. 7). The boundaries of this linear regime can change with promoter strength and NTP concentration; however, even with the strongest promoter tested, ∼6 hours of predictable performance may be achieved with measurable fluorescent protein signal over a wide range of DNA template concentrations. We advocate using the linear regime for cell-free circuit testing or other applications that require linear response. The high DNA concentration (saturation) regime is best suited for applications when maximum yield is desired but the linearity of the DNA–protein relationship is not essential.

**Figure 7:**
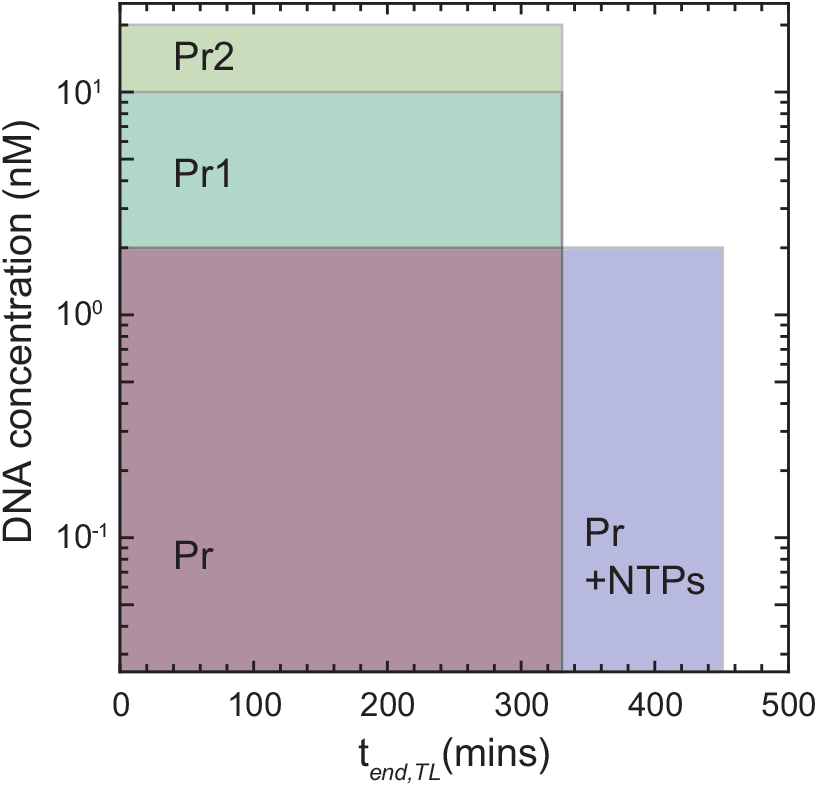
DNA concentration and protein production time boundaries of the linear regime, in which the rate of protein production is approximately constant and the final protein concentration is proportional to the amount of template DNA.

And there are a number of ways in which limits on the capacity of the cell-free breadboard may be raised. The functional lifetime of the system, which in bulk operation is limited by unidentified mechanisms reducing the activity of the transcription/translation machinery, may be increased using dialysis membranes and vesicles, up to 16 and 100 hours, respectively (8, 44). Reaction times may be further extended with the use of microfluidics or other continuous-flow devices, as demonstrated with other cell-free environments (45, 46). Also, the addition of purified proteins such as T7 RNAP or sigma factors could potentially support an increase in capacity at no additional cost to the system; in related work it has been shown that purified GamS protein can be added to prevent degradation of linear DNA (9). Ideally, a combination of strategies should be employed to take maximum advantage of the cell-free breadboard. The ease with which these strategies can be employed, along with the relative simplicity of the system and the control that it offers, makes it a promising platform for synthetic biocircuit prototyping.

## 3 Materials and Methods

### 3.1 **Cell-free system and reactions**

The breadboard environment consists of a crude cytoplasmic extract from *E. coli* containing soluble proteins, including the entire endogenous transcription–translation machinery and mRNA and protein degradation enzymes (7, 8). Detailed instructions for extract preparation can be found in (23). To avoid variation between different extract preparations we used the same batch of extract for all experiments. Reactions took place in 10 *µ*l volumes at 29^◦^C. No significant toxicity was observed for typical deGFP expression experiments when up to 20 *µ*M malachite green dye was included in the reaction; the dye concentration was thus fixed at 10 *µ*M for all experiments. All experiments were run for 14 hours, a time found to be sufficiently long for the translational capacity of the system to be exhausted under all conditions tested.

### 3.2 **Reporters**

Real-time fluorescence monitoring of mRNA dynamics was performed using the malachite green aptamer (MGapt) (13) incorporated in the 3′ UTR of the fluorescent protein reporter genes, 15 bases downstream of the stop codons. This location of MGapt insertion was chosen after a number of other possible locations were tested and found to give less accurate measures of RNA dynamics. For example, incorporation of MGapt within the 5′ UTR upstream of the RBS led to decreased MGapt fluorescence signal, a result that may be due to the preference of 5′ end degradation by the dominant endonuclease in *E. coli*, RNase E (47). This is consistent with a recent study on the Spinach fluorescent RNA aptamer (17) in which it was reported that incorporation of the aptamer in the 3′ UTR region led to stronger fluorescence than in the 5′ UTR. It was also found that a 6-base spacing between the stop codon of deGFP and MGapt affected the protein expression level to some extent, but a 10-base and 15-base spacing showed equivalent MGapt fluorescence signal levels without affecting protein expression. The fluorescent proteins deGFP and deCFP were previously designed to be more translatable in the cell-free system (7). The UTR controlling translation of deGFP (eGFP-Δ_6-229_) and deCFP contained the T7 gene *10* leader sequence for highly efficient translation initiation (7). All transcription units included the T500 transcriptional terminator except for PT7-deGFP-MGapt which contained T7 terminator.

Fluorescence measurements were made in triplicate in a Biotek plate reader at 3 minute intervals using excitation/emission wavelengths set at 610/650 nm (MGapt), 485/525 nm (deGFP), and 433/475 nm (deCFP). Error bars in figures showing fluorescence or integrated fluorescence indicate standard error over replicates. The reported protein production end times are the times at which the protein concentrations reached 95% of their final values, i.e., deGFP(*t_end,TL_*) = 95% × [deGFP]_*end*_.

### 3.3 **Plasmids and Bacterial strains**

Plasmids was created using standard cloning methods. The plasmid pBEST-Luc (Promega) was used as a template for all constructs except for the PT7-deGFP-MGapt construct which was derived from the plasmid pIVEX2.3d (Roche). The same antibiotic resistance gene was used with each plasmid to ensure that any burden on the system due to the expression of these ‘background’ proteins was the same for each construct. All plasmids used are listed in Table S1. Plasmid DNAs used in cell-free experiments were prepared using Qiagen Plasmid Midi prep kits. *E. coli* strains KL740 (which contains lambda repressor to control for Pr promoter) or JM109 were used. LB media with 100 *µ*g/mL carbenicillin was used to culture cells.

The Pr promoter is that of the lambda phage repressor gene *cro*. Promoters Pr1 and Pr2 were each modified from Pr with a single base mutation in the -35 and -10 region, respectively. The sequences, with mutations highlighted by □, are

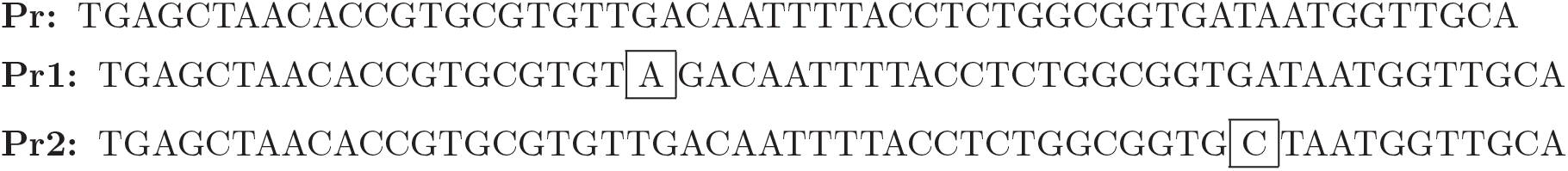

### 3.4 **Preparation of pure mRNA and qRT-PCR**

RNA was transcribed using a linear template PCR-amplified from pIVEX2.3d PT7-deGFP-MGapt including T7 promoter and T7 terminator region. The transcription reaction was prepared as a total volume of 100 *µ*L with 0.1 *µ*M linear DNA template, 20% (v/v) T7 RNA polymerase (Cellscript), 7.5 mM each NTP (Epicentre), 24 mM MgCl_2_ (Sigma), 10% (v/v) 10× transcription buffer, and 1% (v/v) thermostable inorganic pyrophosphatase (New England Biolabs). After an overnight incubation at 37^◦^C, the reaction mixture was run on 1% agarose gel, RNA bands that correpond to full-length transcript were excised and eluted from gel by the Freeze-N-Squeeze column (Biorad) and resuspended in water. Concentrations of purified RNA were determined spectrophotometrically using Nanodrop.

For qRT-PCR, 1 *µ*L samples were taken at different time points from a tube containing reaction mixture at 29^◦^C and diluted 50-fold in water. These samples were stored at -80^◦^C until used. Afterward the samples were further diluted to a final dilution of 1:5000. Two *µ*L of samples were analyzed in 50 *µ*L reactions of the Power SYBR Green RNA-to-CT 1-Step kit (Life Technologies) in the MX3005 real-time PCR machine (Agilent Technologies). Primers amplified a region of the deGFP gene closer to its 3′ end (424-597 nt) and were used at 0.3 *µ*M concentrations. Concentrations of deGFP-MGapt RNA in the sample were determined from a standard curve of dilutions of purified mRNA in a range from 0.6 to 60 pM mRNA per PCR reaction.

## Acknowledgement

The authors thank Eduardo Sontag and members of the Murray Group for useful discussions, and S. C. Livingston, P. Rovo, G. Smith for helpful comments on the manuscript. This research is funded in part by the Gordon and Betty Moore Foundation through Grant GBMF2809 to the Caltech Programmable Molecular Technology Initiative, and by the Defense Advanced Research Projects Agency (DARPA/MTO) Living Foundries program, contract number HR0011-12-C-0065. Z.A.T. was partially supported by grants TAMOP-4.2.1-B-11/2/KMR-2011-0002, TAMOP-4.2.2./B-10/1-2010-0014 and OTKA NF 104706.

The views and conclusions contained in this document are those of the authors and should not be interpreted as representing official policies, either expressly or implied, of the Defense Advanced Research Projects Agency or the U.S. Government.

## Supporting Information Available

This material is available free of charge via the Internet at http://pubs.acs.org/.

## Graphical TOC Entry

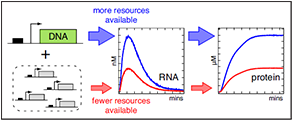

## Supporting Information

### Alternative performance metrics

For the purposes of characterization of a particular biocircuit testing environment, *in vitro* or *in vivo*, there are any number of performance metrics that may be used. We have chosen to use integrated mRNA and final protein concentration since they intuitively represent the total transcriptional and translational capacity of a system. Other metrics, such as the mRNA and protein production rates and end times, are complementary to those used in this work and may be particularly informative depending on the specific circuit or system requirement.

In Figs. S16 and S17 we show the maximum deGFP and MGapt production rates as a function of reporter concentration under different conditions. Using these measures, stark differences between simple one-stage gene expression and expression from the two-stage T7 cascade can be seen; for example, at low reporter concentrations, the maximum production rate of deGFP in the cascade is considerably larger than the one-stage rate, even when the strongest promoter is used. The cascade deGFP rates are also relatively flat with respect to reporter concentration. There is also a clear effect of NTPs on peak rates when a cascade is used versus simple expression. In the former case, additional NTPs provide for a significant increase in the maximum protein and mRNA production rates, whereas NTPs have little to no effect on maximum production rates for the Pr-deGFP-MGapt construct.

The deGFP production end time *t_end,TL_* is shown for all conditions in Fig. S18. *t_end,TL_* is between 330 and 350 minutes for simple expression in the ‘linear’ regime, but as high as ∼580 minutes in some of the conditions and concentrations tested. The end time is a particularly good measure of system capacity when extended performance is required.

## Supplementary Figures

**Figure S1:**
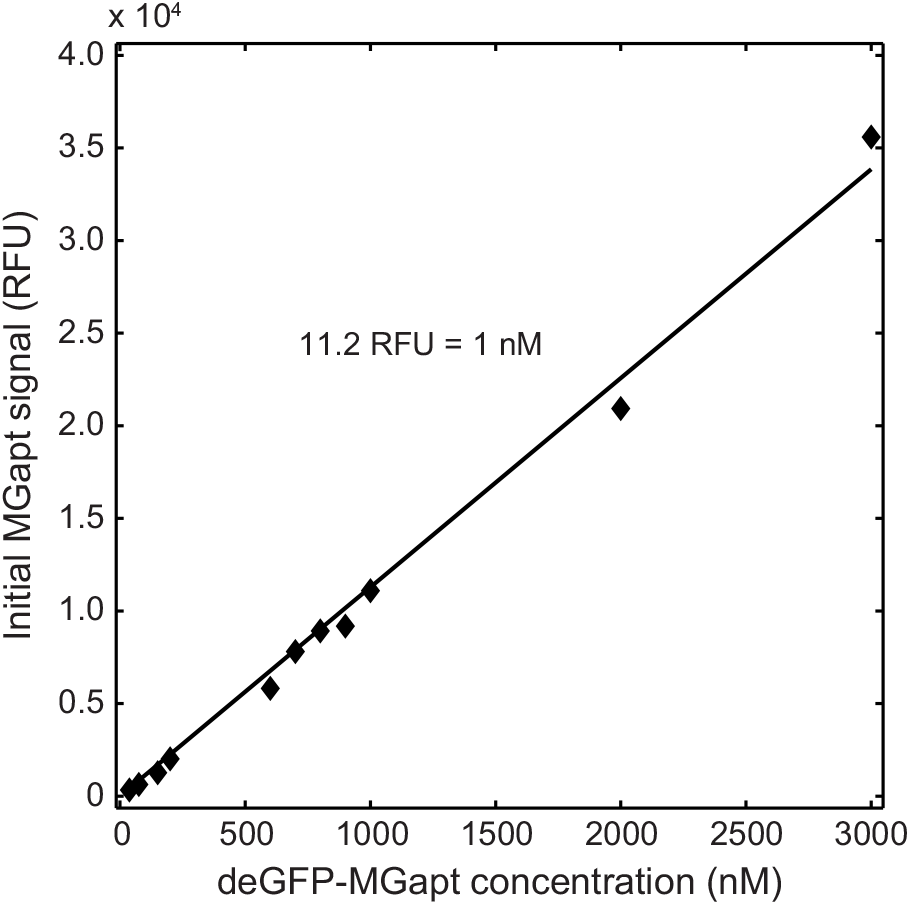
Calibration curve relating concentration of purified deGFP-MGapt transcript to initial MGapt fluorescence signal.

**Figure S2:**
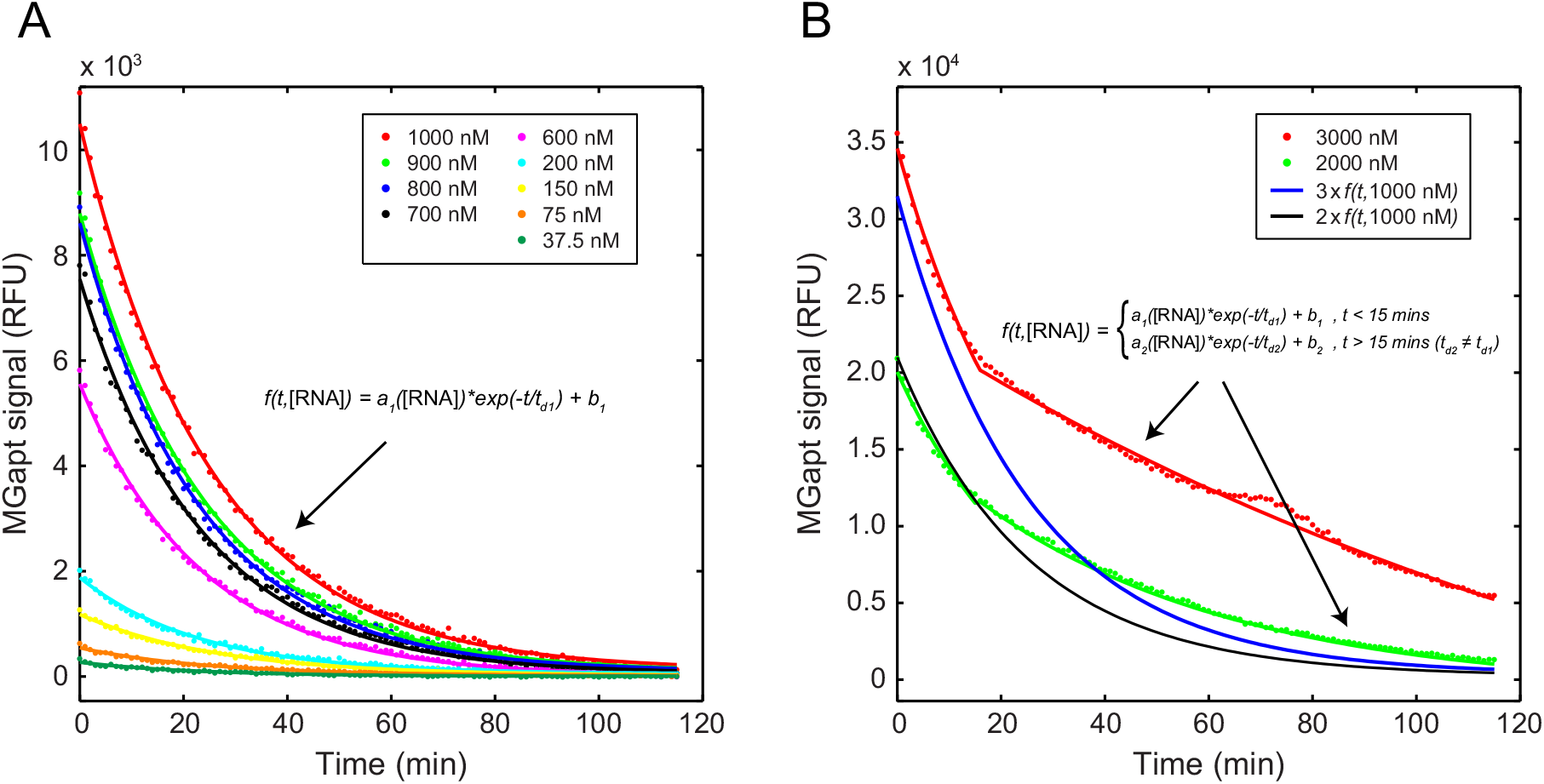
Decay of purified deGFP-MGapt transcripts in the cell-free breadboard. (A) Degradation is well-described by single exponential decay with 16–18 min half-lives for a wide range of concentrations. (B) At high concentrations, the decay curves follow those of the lower concentrations only to 15 min, after which the half-lives increase dramatically.

**Figure S3:**
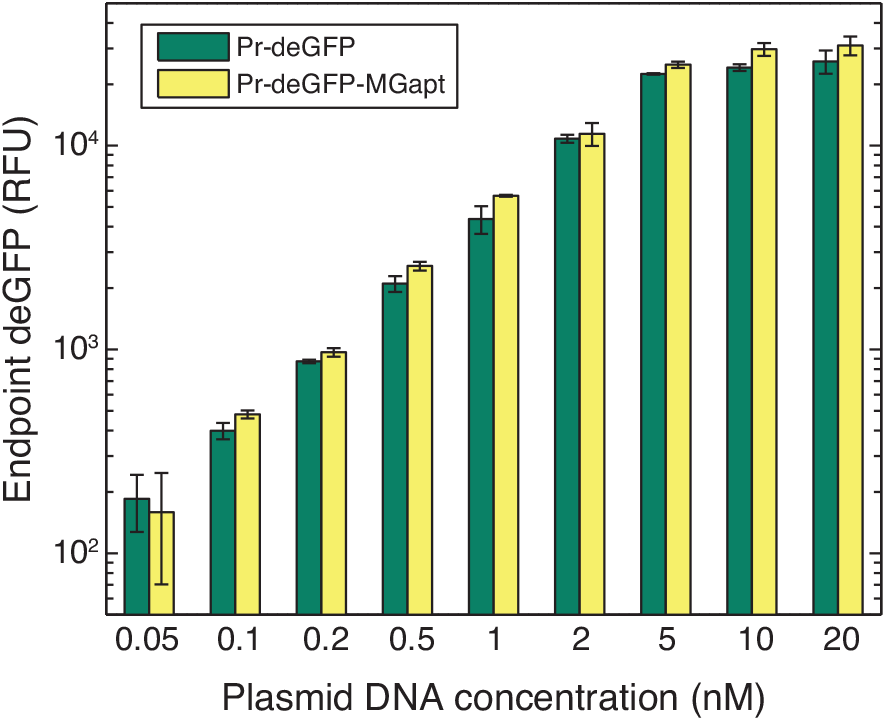
Comparison of total deGFP fluorescence produced by Pr-deGFP and Pr-deGFP-MGapt constructs. The incorporation of MGapt in the 3′ UTR downstream of deGFP leads to only a slight increase in the amount of protein produced relative to deGFP alone, a result we attribute to an increase in the stability of the fusion transcript conferred by the MGapt.

**Figure S4:**
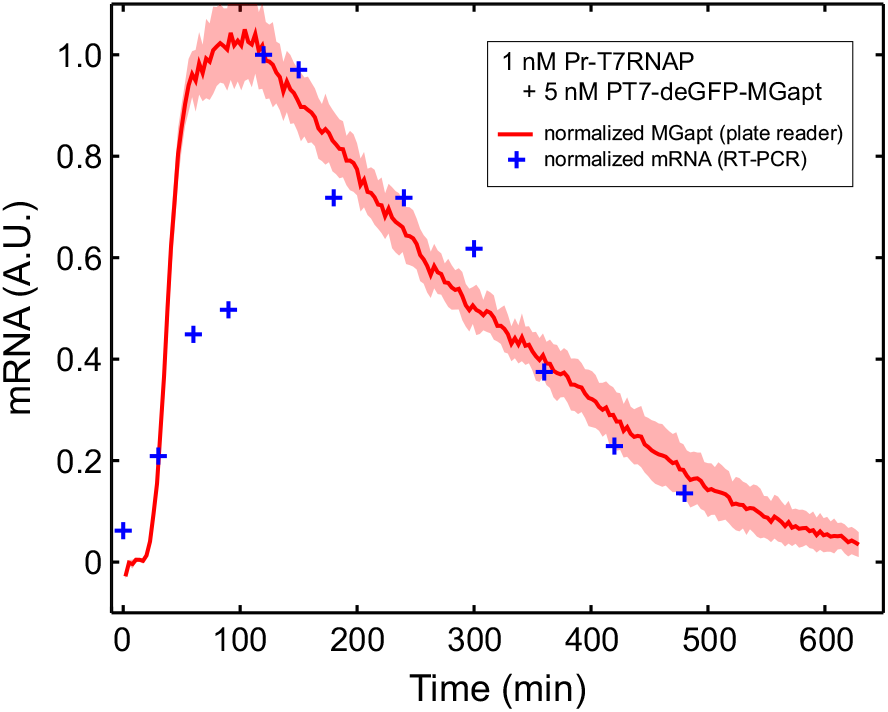
Comparison of normalized real-time PCR measurements and MGapt signal for the T7 RNAP cascade circuit (described in the section titled “Performance of a simple transcription–translation cascade”) with 1 nM Pr-T7 RNAP and 5 nM PT7-deGFP-MGapt. Shaded region indicates standard error over replicates.

**Figure S5.**
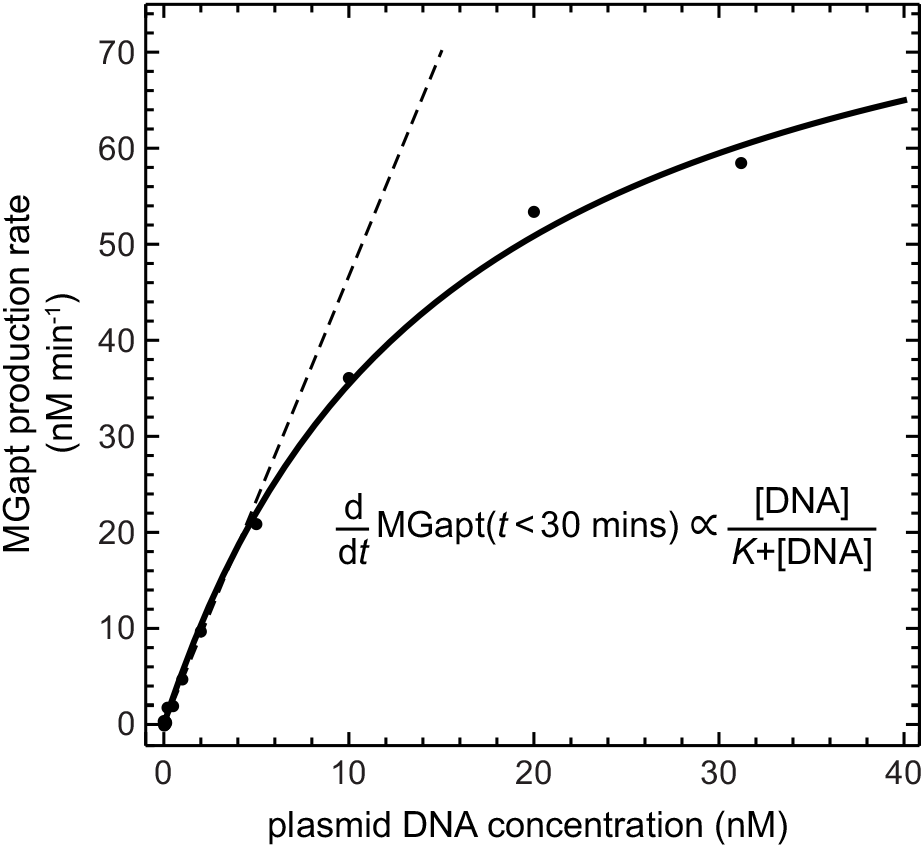
MGapt production at early times suggest transcriptional machinery is saturated as DNA concentration increases. Fits of MGapt production rates in the first 30 minutes of expression follow a Michaelis-Menten form with Michaelis constant *K* ∼15 nM.

**Figure S6:**
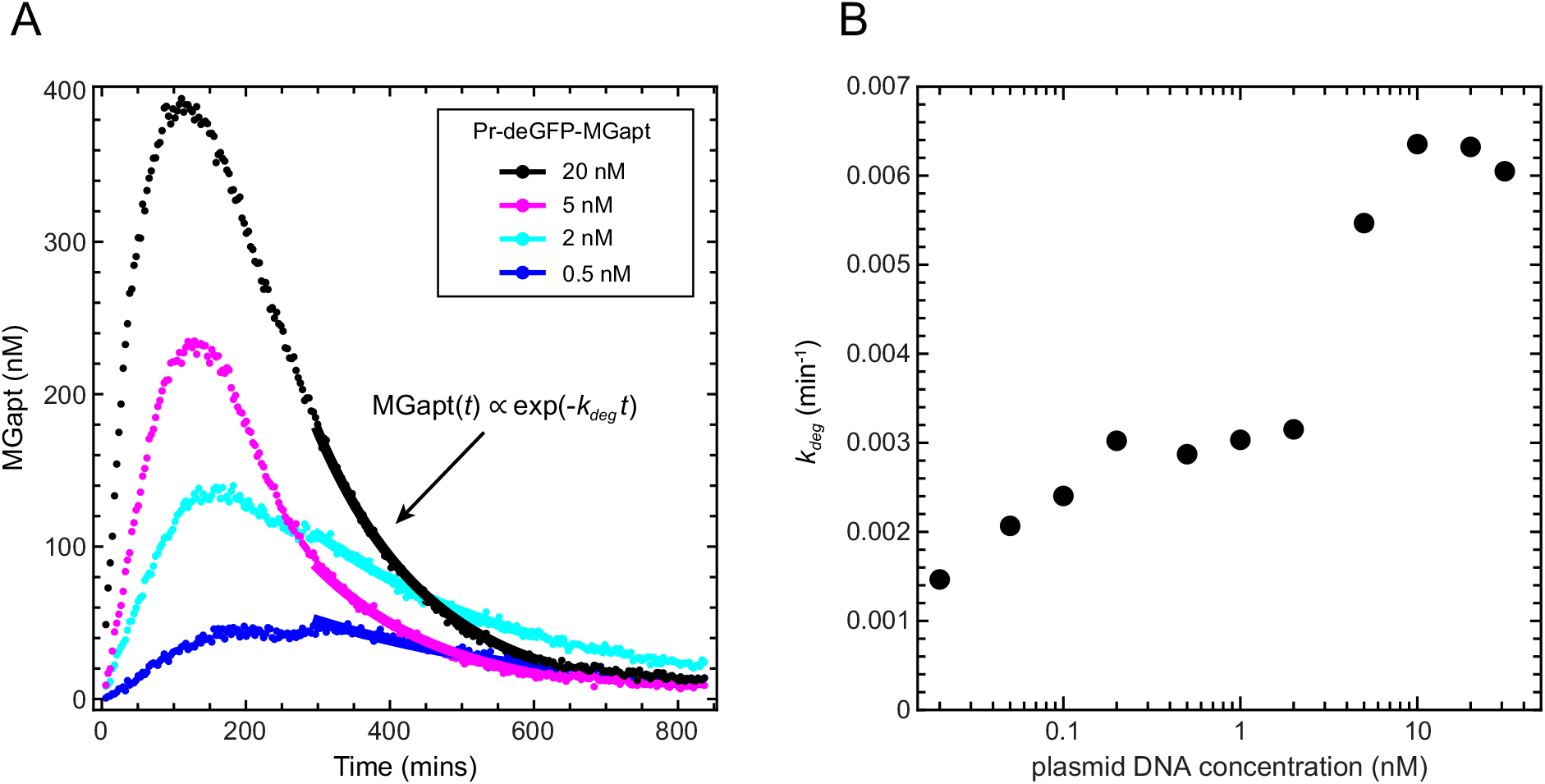
(A) MGapt expression curves are well-fit by a single exponential at late times. (B) Decay constants *k_deg_* determined using a single exponential fit to data after 5 hours of expression. *k_deg_* increases with increasing DNA concentration.

**Figure S7:**
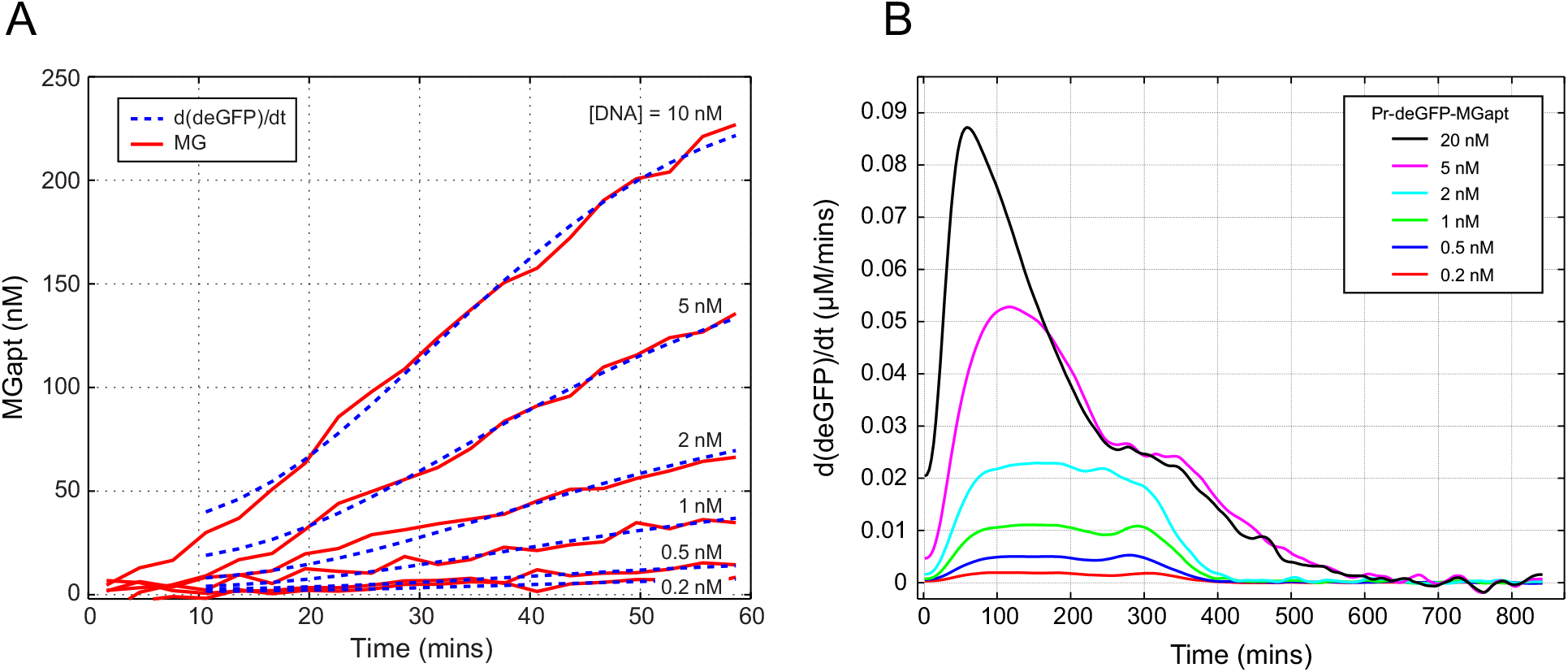
(A) The time derivative of the (spline fit) deGFP concentration is proportional to the MGapt concentration in the first hour of expression. (B) As the experiment goes on and the behavior of the system is no longer ideal, the shapes of the 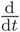deMGapt(*t*) deviate from MGapt(*t*) (although there remain significant qualitative similarities between them; cf. Fig. 1C.

**Figure S8:**
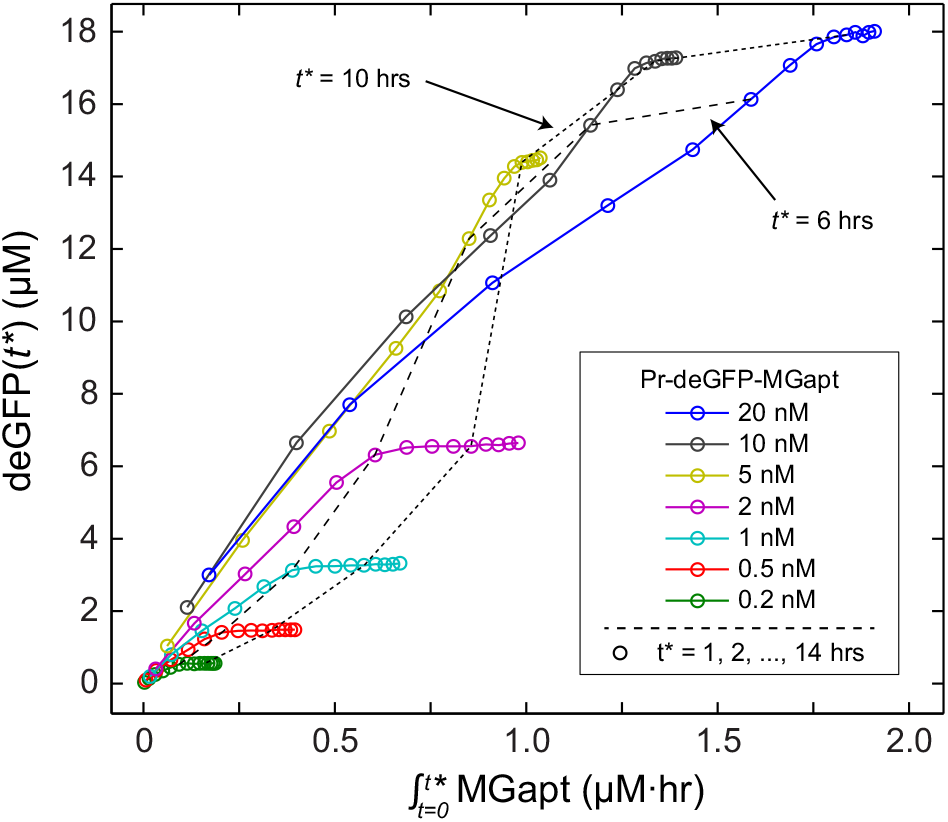
deGFP at various times *t*\* versus MGapt level integrated from time *t* = 0 to *t* = *t*\* for a range of Pr-deGFP-MGapt concentrations, with *t*\* = 1, 2, …, 14 hrs indicated with ○. Below the ‘linear’–‘saturation’ regime transition concentration, 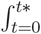MGapt appears proportional to deGFP(*t\**) until protein synthesis stops at *t*\* ≈ 340 minutes, albeit with a proportionality constant that is different for different DNA template concentrations. Above the transition concentration, protein production stops later in the experiment and the 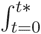MGapt plateau is relatively short. Dashed lines showing *t*\* = 6 hrs and *t*\* = 10 are shown for reference.

**Figure S9:**
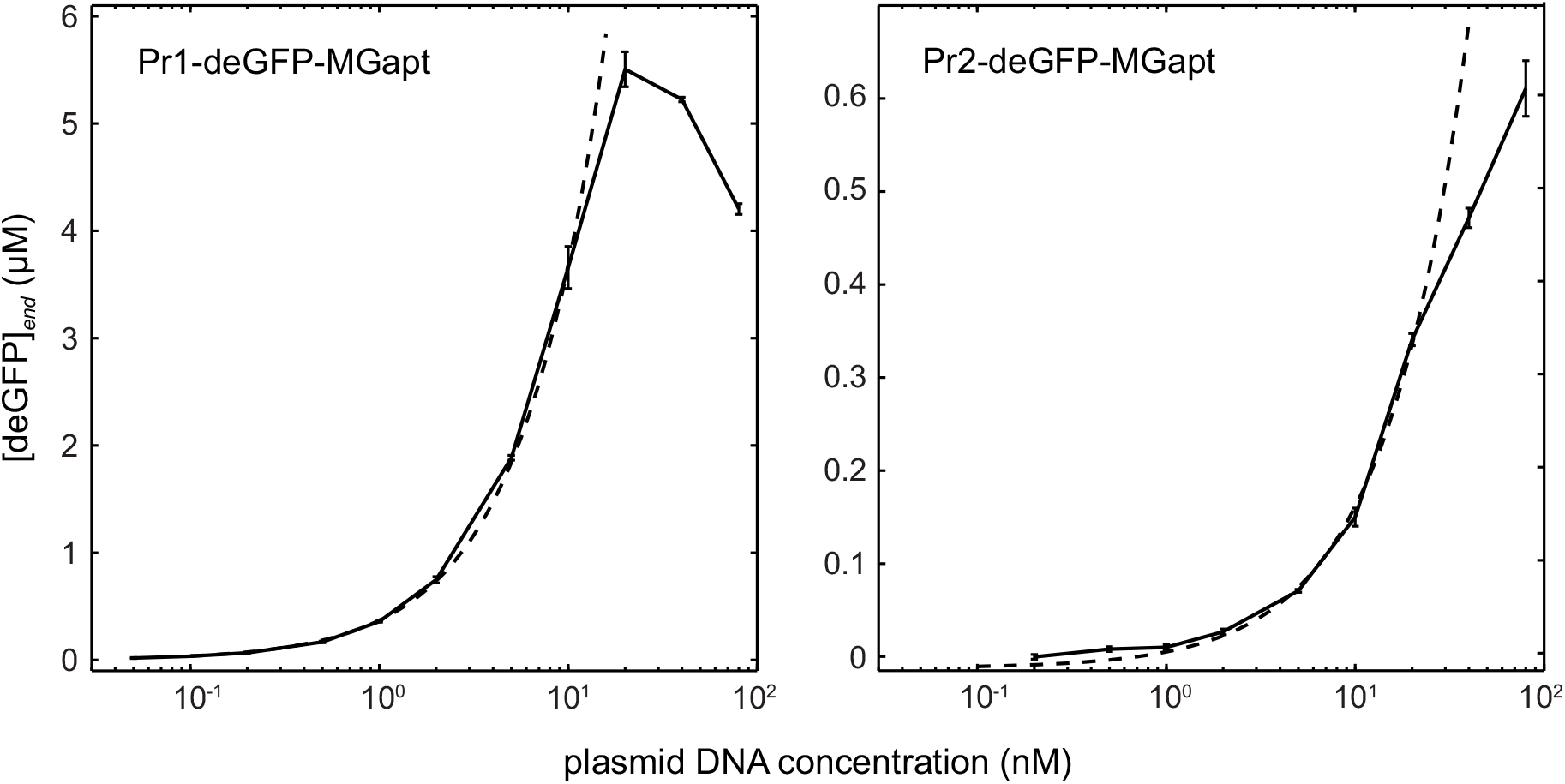
Endpoint deGFP versus plasmid concentration for weak promoters Pr1 (left) and Pr2 (right). Unlike the Pr promoter for which the linear regime ends at 2–5 nM DNA, for Pr1 and Pr2, the linear regime extends up to ∼10 nM and ∼20 nM, respectively. Dashed line shows a linear fit to the data.

**Figure S10:**
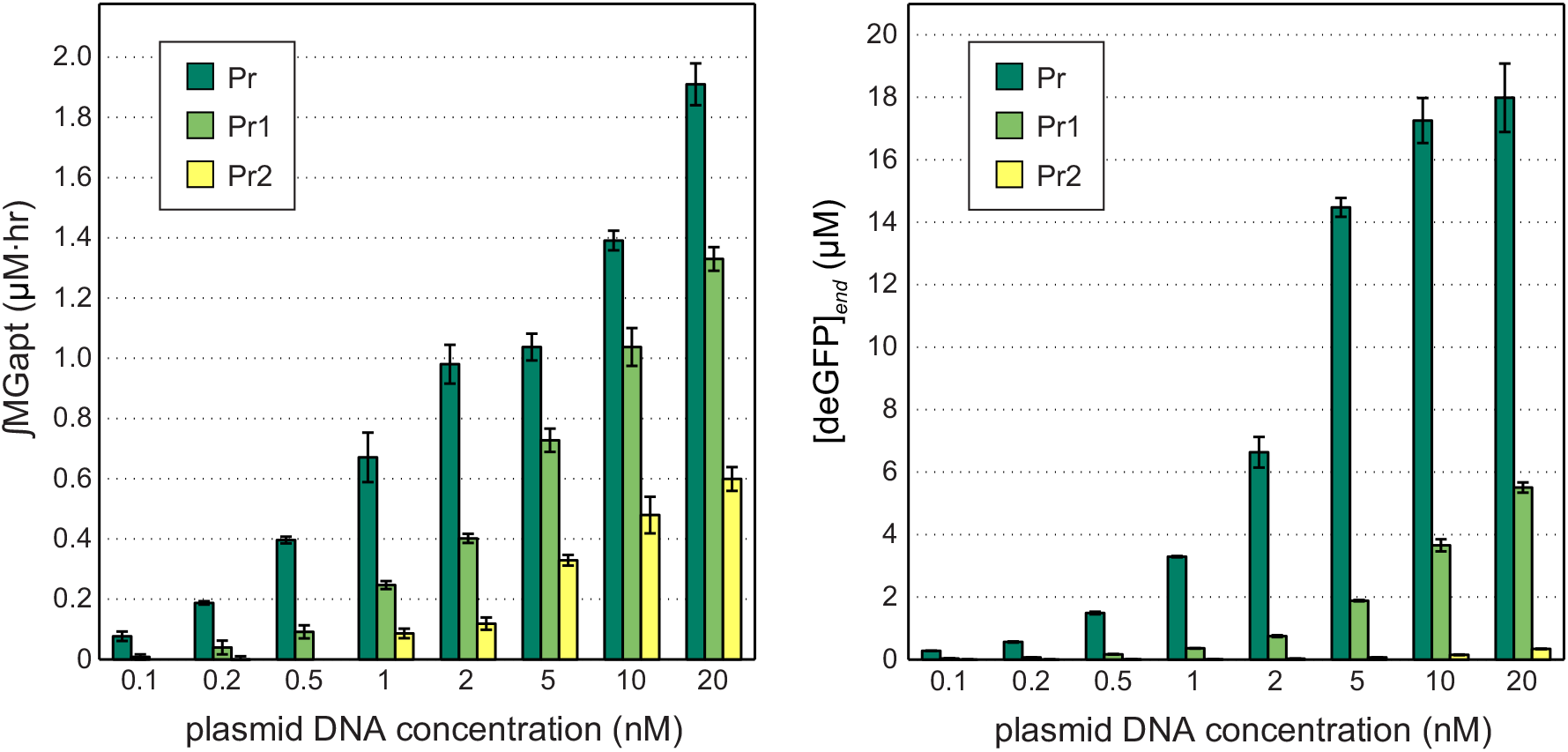
Comparison of integrated MGapt (left) and endpoint deGFP (right) for promoters Pr, Pr1, and Pr2.

**Figure S11:**
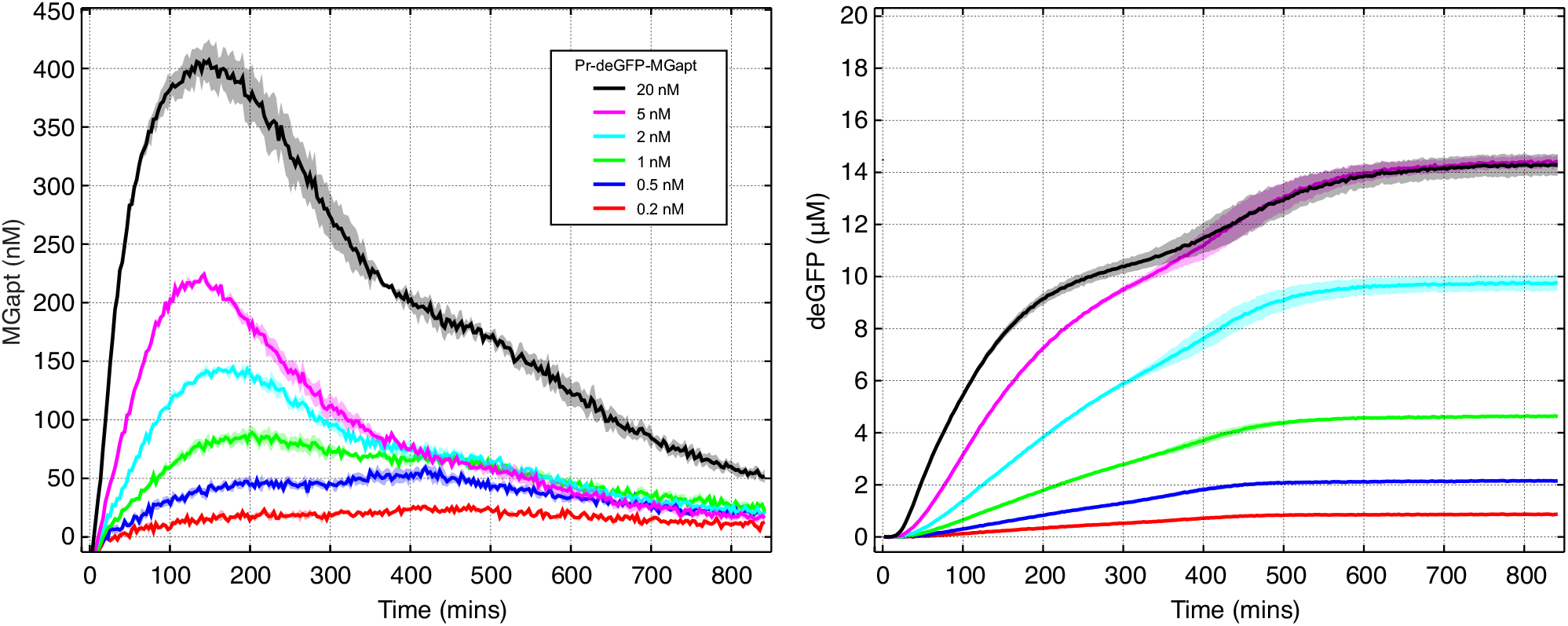
MGapt (left) and deGFP (right) expression kinetics when breadboard is supplemented with 1.25 mM of each of the four NTPs. Shaded regions indicate standard error over replicates.

**Figure S12:**
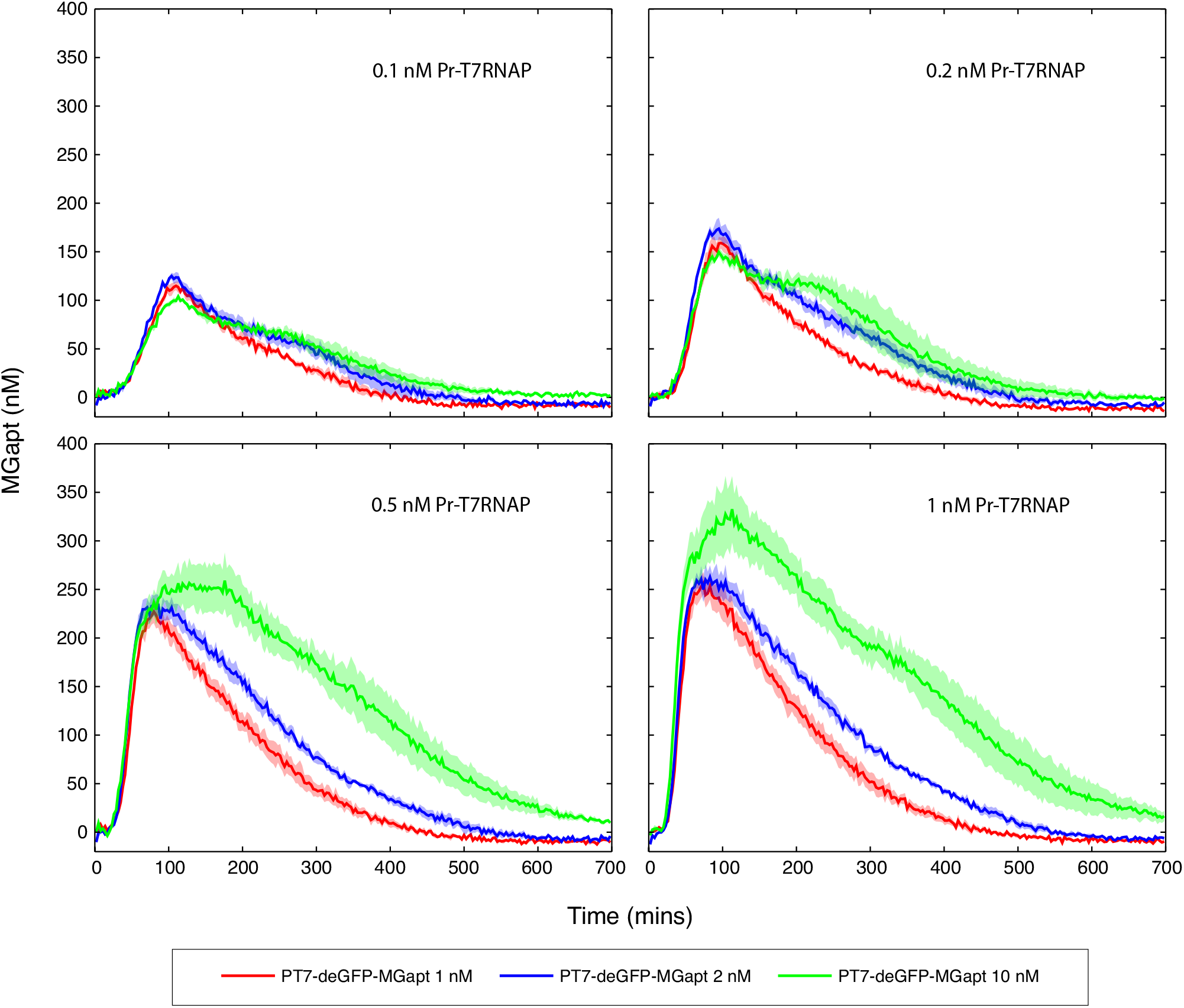
MGapt expression curves for T7 cascade tested with four different concentrations of first-stage T7 RNAP plasmid (0.1, 0.2, 0.5, and 1 nM Pr-T7 RNAP) and three different concentrations of the second-stage plasmid (1, 2, and 10 nM PT7-deGFP-MGapt). Shaded regions indicate standard error over replicates.

**Figure S13:**
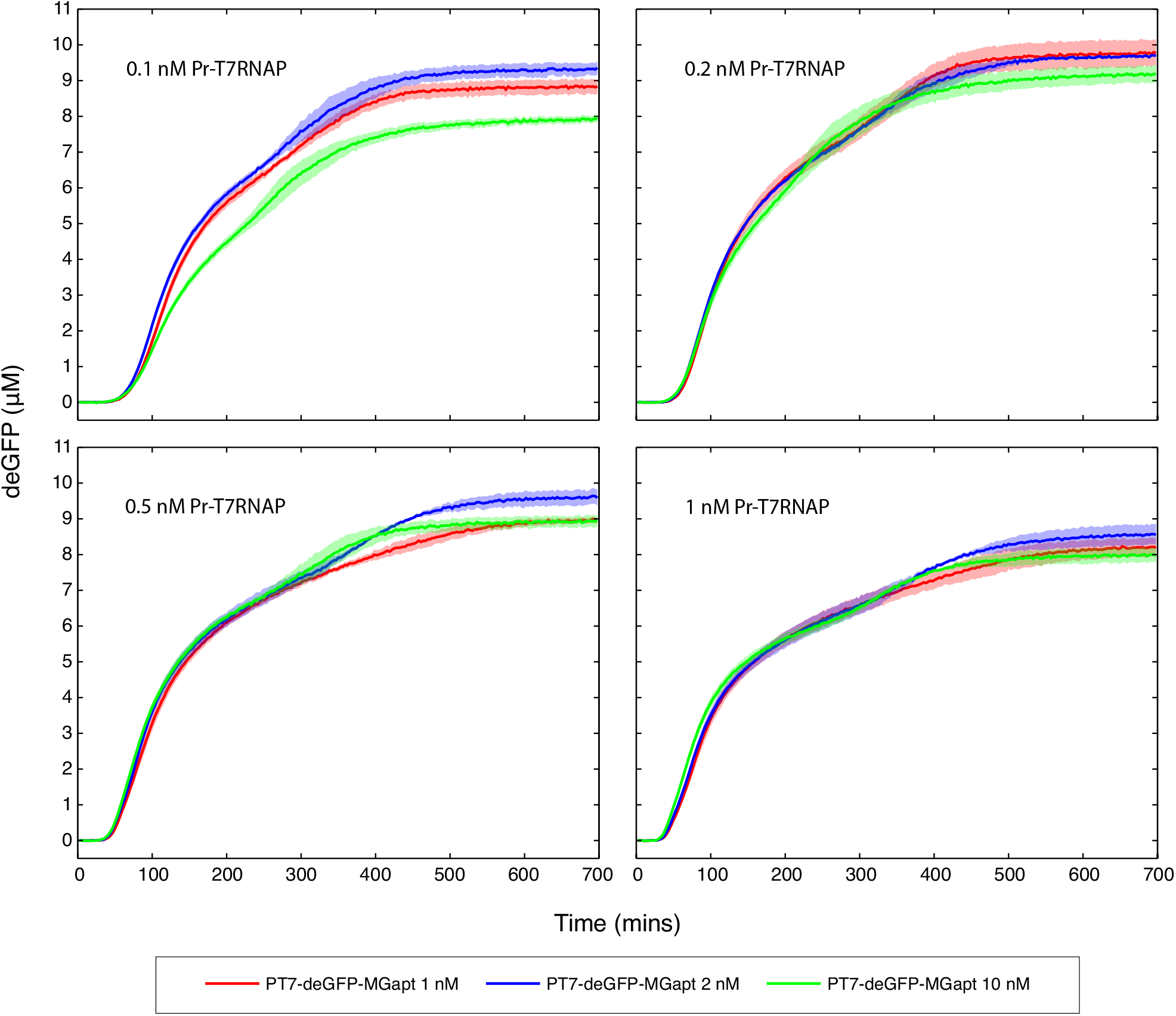
deGFP expression curves for T7 cascade tested with four different concentrations of the first-stage T7 RNAP plasmid (0.1, 0.2, 0.5, and 1 nM Pr-T7 RNAP) and three different concentrations of the second-stage plasmid (1, 2, and 10 nM PT7-deGFP-MGapt). Shaded regions indicate standard error over replicates.

**Figure S14:**
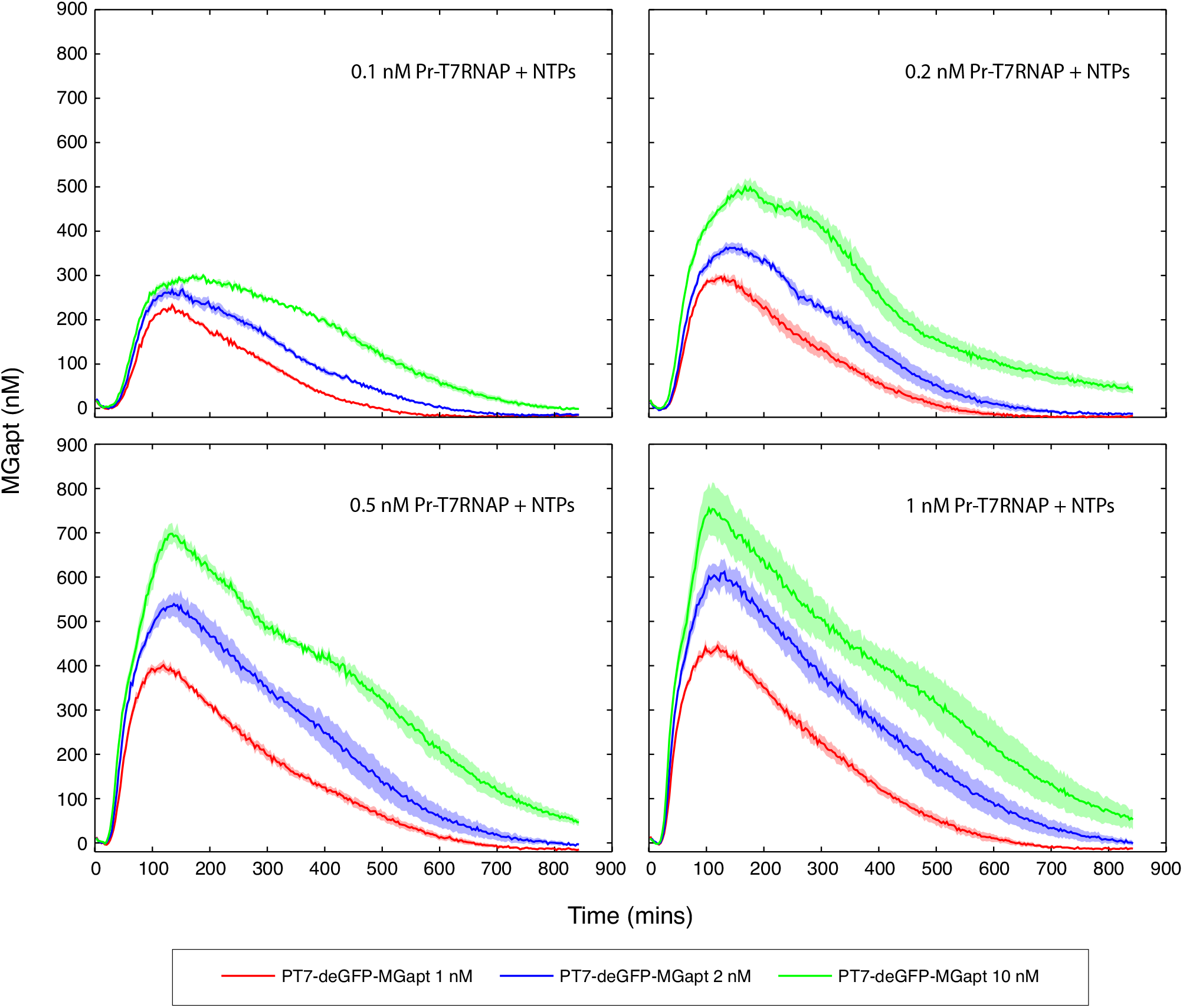
MGapt expression curves for T7 cascade tested with four different concentrations of first-stage T7 RNAP plasmid (0.1, 0.2, 0.5, and 1 nM Pr-T7 RNAP), three different concentrations of the second-stage plasmid (1, 2, and 10 nM PT7-deGFP-MGapt), and with the cell-free breadboard supplemented with 1.25 mM of each of the four NTPs. Shaded regions indicate standard error over replicates.

**Figure S15:**
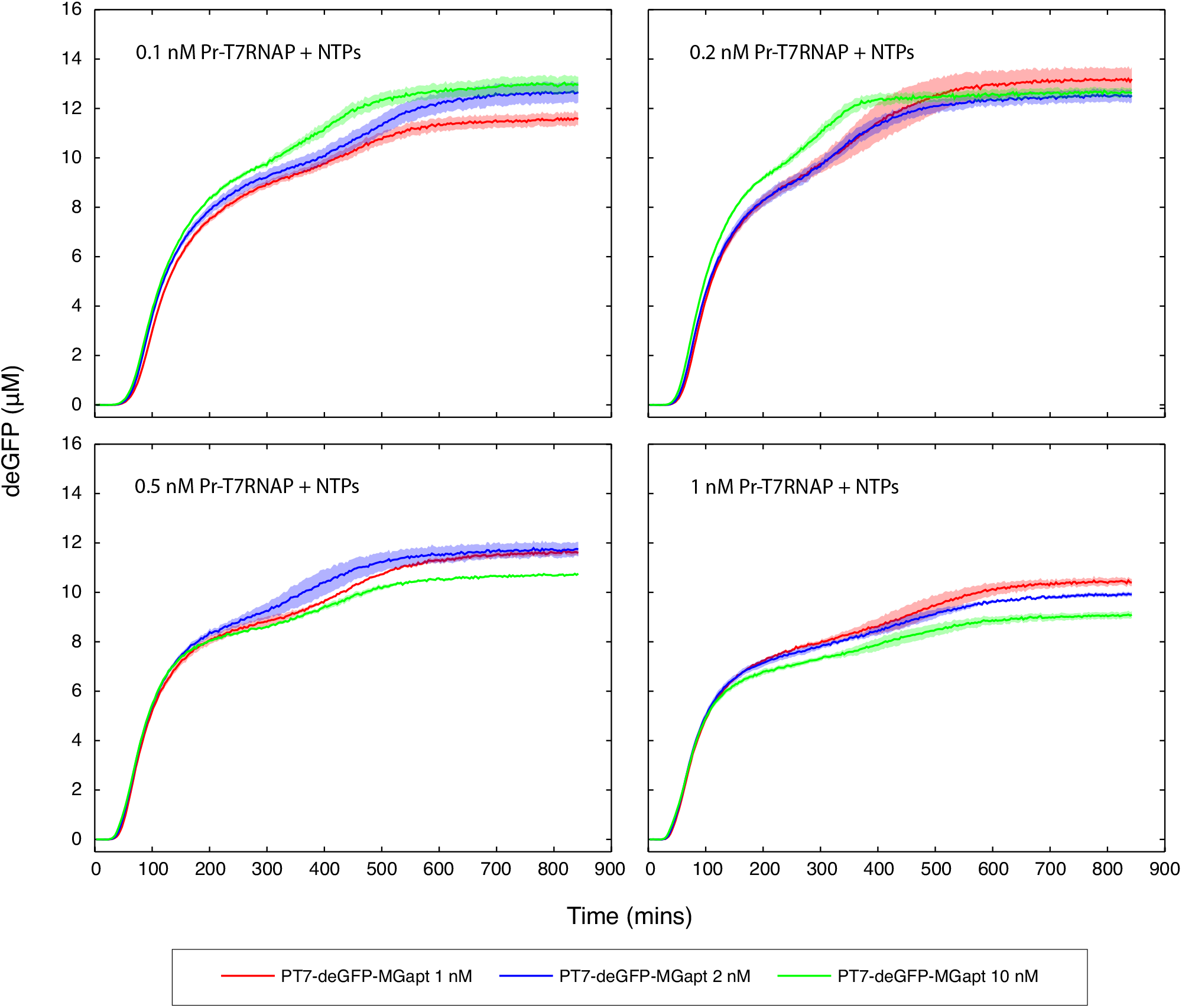
deGFP expression curves for T7 cascade tested with four different concentrations of the first-stage T7 RNAP plasmid (0.1, 0.2, 0.5, and 1 nM Pr-T7 RNAP), three different concentrations of the second-stage plasmid (1, 2, and 10 nM PT7-deGFP-MGapt), and with the cell-free breadboard supplemented with 1.25 mM of each of the four NTPs. Shaded regions indicate standard error over replicates.

**Figure S16:**
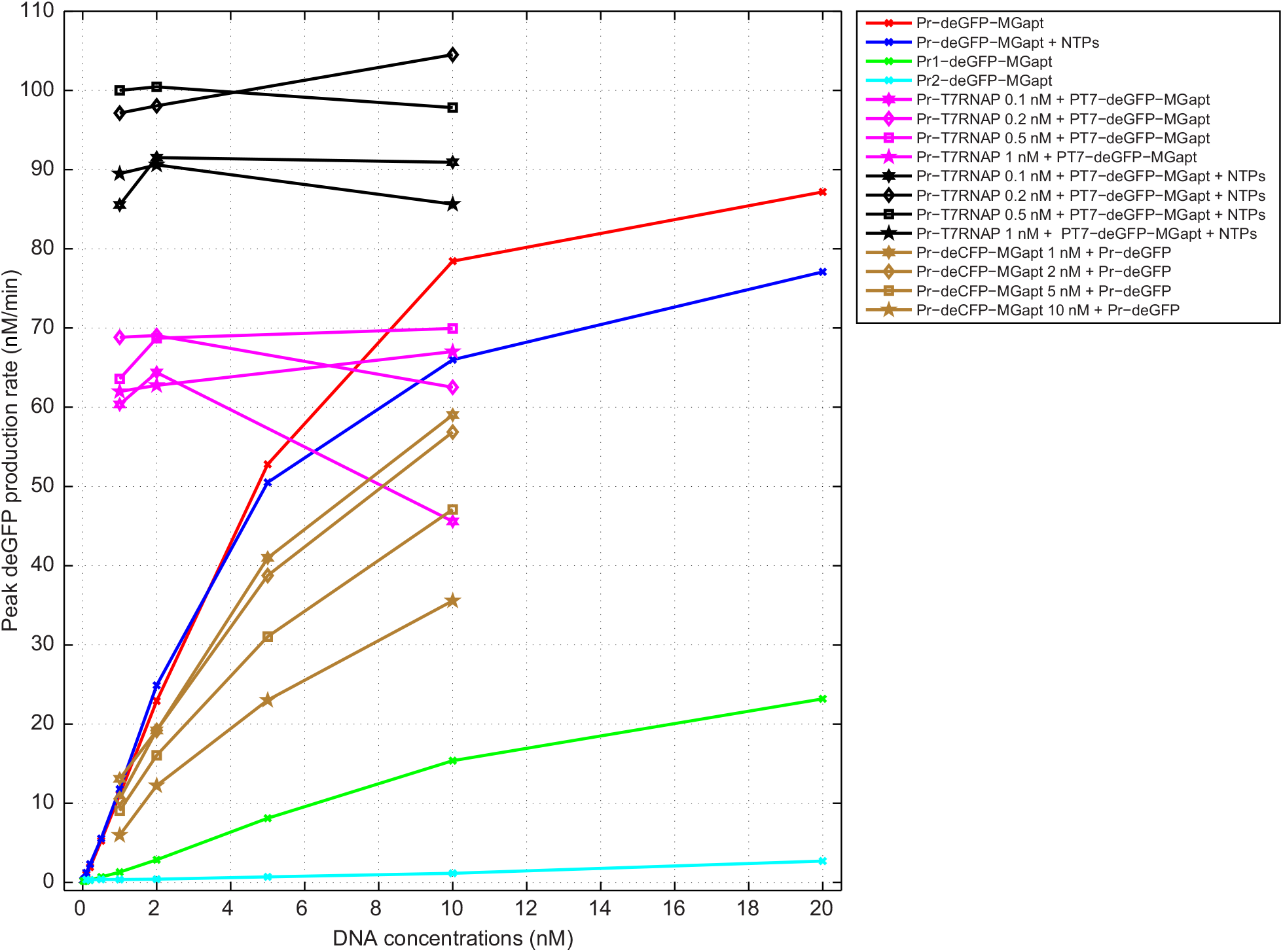
Maximum deGFP production rate as a function of reporter concentration under different conditions.

**Figure S17:**
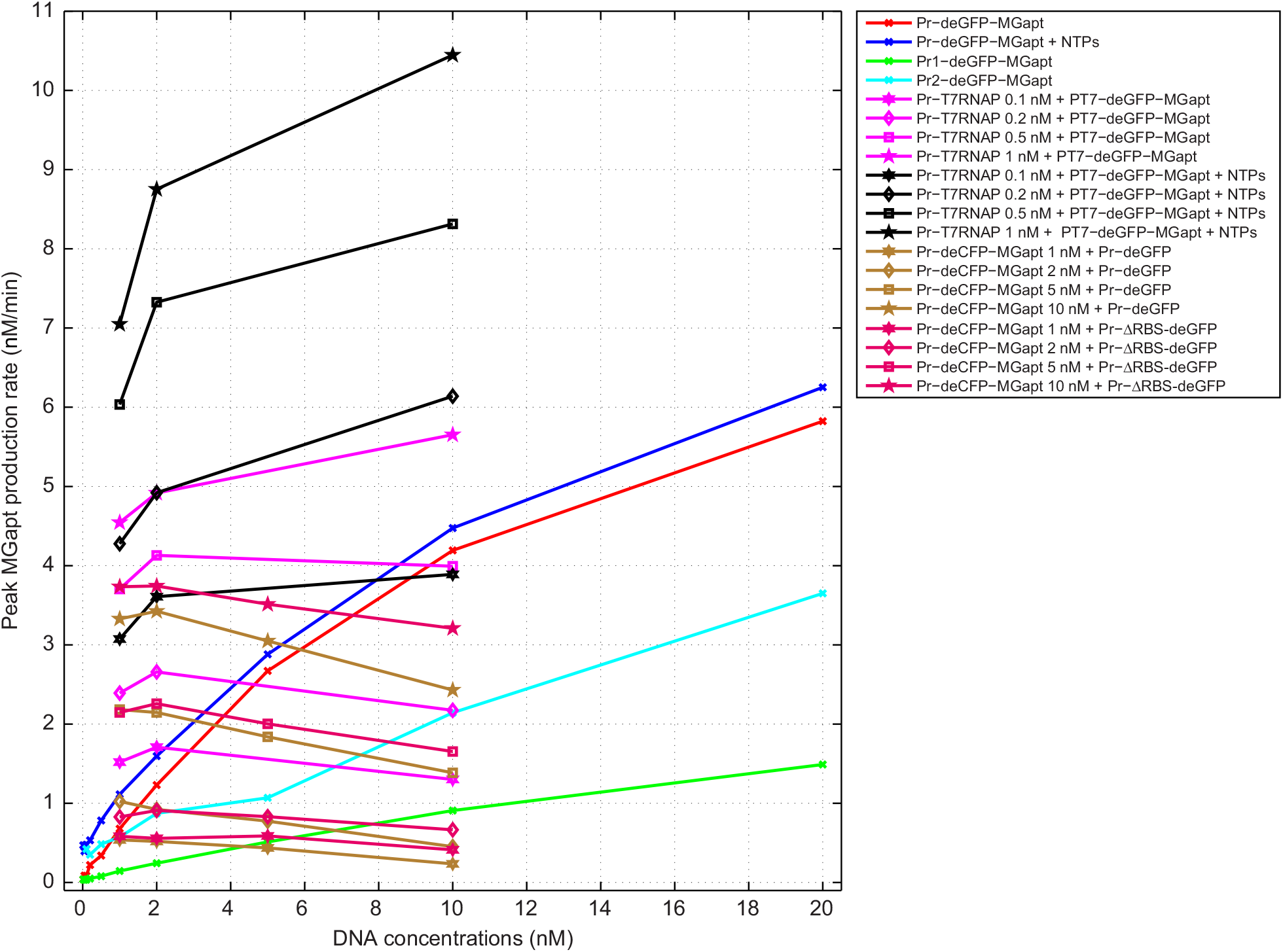
Maximum MGapt production rate as a function of reporter concentration under different conditions.

**Figure S18:**
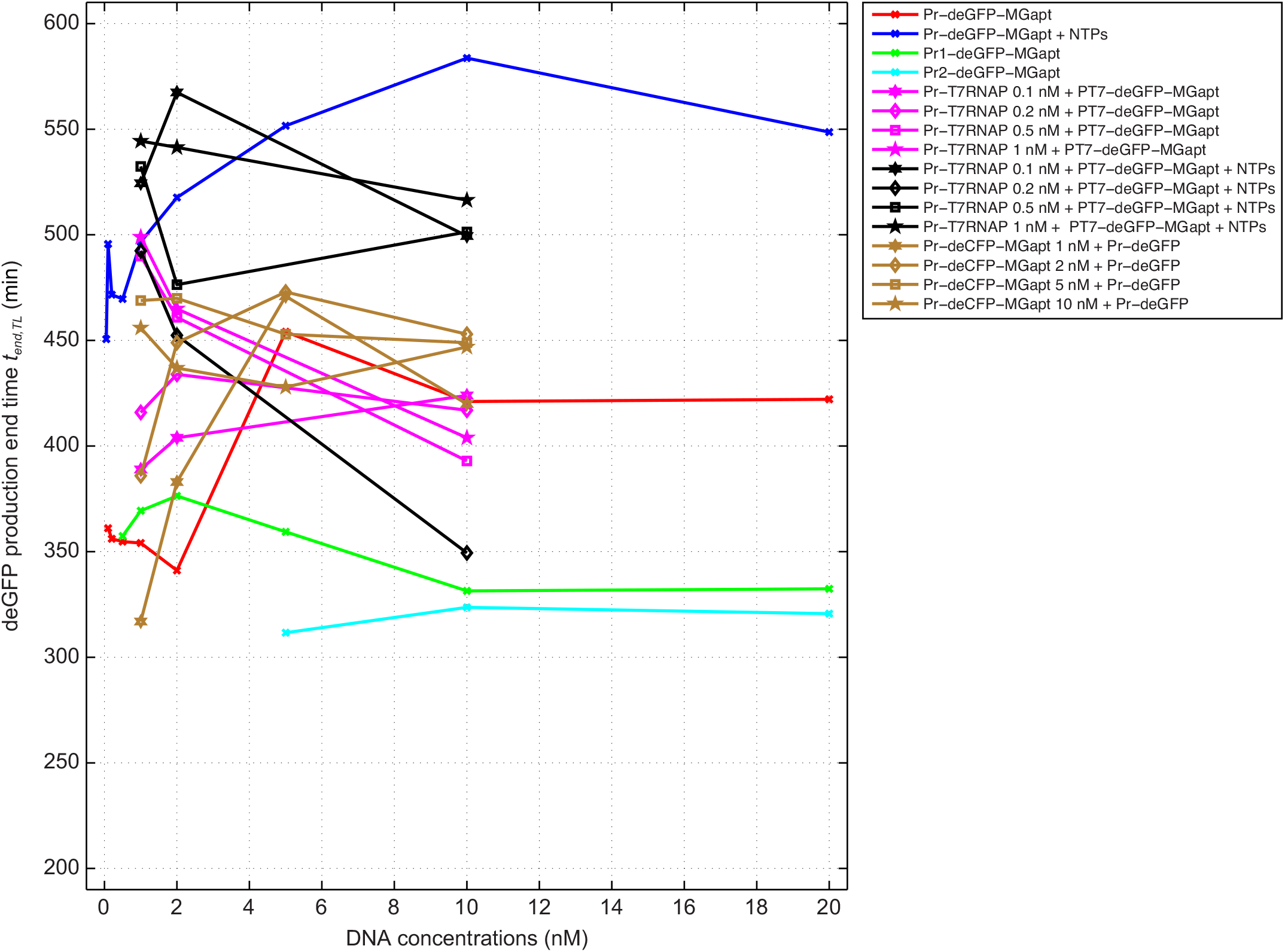
deGFP production end time *t_end,TL_* under different conditions.

## Supplementary tables

**Table S1.** Genotypes of the plasmids used in this study.

| Plasmid name | Transcription unit (TU) | TU size (bp) | Backbone/resistance |
| --- | --- | --- | --- |
| pBEST-Pr-GFP | P<sub>R</sub>:deGFP:T500 | 810 | ColE1/Amp<sup>R</sup> |
| pBEST-Pr-ΔRBS-GFP | P<sub>R</sub>:ΔRBS-deGFP:T500 | 775 | ColE1/Amp<sup>R</sup> |
| pBEST-Pr-GFP-MG | P<sub>R</sub>:deGFP-MGapt:T500 | 869 | ColE1/Amp<sup>R</sup> |
| pBEST-Pr1-GFP-MG | P<sub>R1</sub>:deGFP-MGapt:T500 | 869 | ColE1/Amp<sup>R</sup> |
| pBEST-Pr2-GFP-MG | P<sub>R2</sub>:deGFP-MGapt:T500 | 869 | ColE1/Amp<sup>R</sup> |
| pBEST-Pr-CFP-MG | P<sub>R</sub>:deCFP-MGapt:T500 | 869 | ColE1/Amp<sup>R</sup> |
| pBEST-Pr-T7RNAP | P<sub>R</sub>:T7 RNAP:T500 | 3032 | ColE1/Amp<sup>R</sup> |
| pBEST-Pr1-T7RNAP | P<sub>R1</sub>:T7 RNAP:T500 | 3032 | ColE1/Amp<sup>R</sup> |
| pBEST-Pr2-T7RNAP | P<sub>R2</sub>:T7 RNAP:T500 | 3032 | ColE1/Amp<sup>R</sup> |
| pIVEX-pT7-GFP-MG | P<sub>T7</sub>:deGFP-MGapt:T7term | 939 | ColE1/Amp<sup>R</sup> |

